# Variations of floral temperature in changing weather conditions

**DOI:** 10.1101/2024.03.04.583410

**Authors:** Michael J. M. Harrap, Natasha de Vere, Natalie Hempel de Ibarra, Heather M. Whitney, Sean A. Rands

## Abstract

1. Floral temperature is a flower characteristic that has the potential to impact the fitness of flowering plants and their pollinators. Likewise, the presence of floral temperature patterns, areas of contrasting temperature across the flower, can have similar impacts on the fitness of both mutualists.
2. It is currently poorly understood how floral temperature changes under the influence of different weather conditions, and how floral traits may moderate these changes. Such weather dependency will impact how stable floral temperatures are over time and their utility to plant and pollinator. The stability of floral temperature cues is likely to facilitate effective plant-pollinator interactions and play a role in the plant’s reproductive success.
3. We use thermal imaging to monitor how floral temperatures and temperature patterns of four plant species (*Cistus* ‘snow fire’ and ‘snow white’, *Coreopsis verticillata* and *Geranium psilostemon*) change with several weather variables (illumination, temperature; windspeed; cloud cover; humidity and pressure) during times that pollinators are active.
4. All weather variables influenced floral temperature in one or more species. The directionality of these relationships were similar across species. In all species light conditions (illumination) had the greatest influence on floral temperature overall, and in generation of contrasting temperatures between parts of the flower, temperature patterns. The effect sizes of other weather variables were lower and more varied across the four species. Most likely, floral traits such as pigmentation and structure influence these relationships between weather conditions and generation of floral temperature.
5. Synthesis: Floral temperature and the extent to which flowers showed contrasting temperature patterns were influenced predominantly by light conditions. However, several weather variables had additional, lesser, influences. Furthermore, differences in floral traits, pigmentation and structure, likely resulted in differences in temperature responses to given conditions between species and different parts of the same flower. However, floral temperatures and contrasting temperature patterns that are sufficiently elevated for detection by pollinators were maintained across most conditions if flowers received moderate illumination. This suggests the presence of elevated floral temperature and contrasting temperature patterns are fairly constant and may have potential to influence plant-pollinator interactions across weather conditions.

## Introduction

Floral temperature has important influences on plant biology (van der Kooi et al., 2019). It can affect flower metabolic processes (Borghi et al., 2019, 2017; Borghi and Fernie, 2017), development (Karlsson et al., 1989; Rodrigo and Herrero, 2002), pollen, ovule and seed viability (Hinojosa et al., 2019; Mu et al., 2017) and plant water balance (Corbet et al., 1979a, 1979b; Gates, 1968). The temperatures of floral surfaces, such as petals and reproductive structures, also influence how floral visitors respond to flowers. Differences in the temperature of the floral surfaces between flowers can influence pollinator foraging choices, acting as part of the flower’s multimodal display (Raguso, 2004; Leonard et al., 2012; Whitney et al., 2008; Hammer et al., 2009; Rands et al., 2023). Bees show unlearnt preferences for flowers with elevated temperatures (Dyer et al., 2006; Whitney et al., 2008). Elevated floral temperatures help insect visitors to maintain temperature thresholds for flight, and reduce the amount of energy these visitors need to warm themselves up during foraging (Herrera, 1995; Heinrich, 1979, 2004; Rands and Whitney, 2008; Seymour et al., 2003b; Seymour and Matthews, 2006). Elevated floral temperature thus effectively acts as an additional floral reward. However, particularly high floral surface temperatures can become a deterrent to visitors (Norgate et al., 2010; Shrestha et al., 2018). Additionally, bees can learn to distinguish hotter and colder flowers that differ in rewards (Whitney et al., 2008). Furthermore, flowers frequently show regions of contrasting temperature across their surfaces, a ‘temperature pattern’ (Atamian et al., 2016; Dietrich and Körner, 2014; Rejšková et al., 2010). Bumblebees can learn to distinguish flowers based on differences in the shape or arrangement of floral temperature patterns (Harrap et al., 2017, 2019) and use these to learn the location of rewards within flowers, improving flower handling (Harrap et al., 2020b).

Temperature perception in insects is common (Altner and Loftus, 1985; Nishikawa et al., 1992; Steinbach and Gottsberger, 1995), and similar foraging responses to floral temperature cues have been seen across a wide range of insect pollinators (Angioy et al., 2004; Atamian et al., 2016). Nevertheless, it is reasonable to assume that the level of temperature difference required to elicit pollinator foraging responses is likely to vary with pollinator species. While insect electrophysiology recordings (Altner and Loftus, 1985; Lacher, 1964; Ruchty et al., 2010) and experiments with restrained bees (Hammer et al., 2009) have seen responses to small surface temperature differences (<1°C), larger differences (>2°C) in temperature between flowers seem to be required to elicit changes in pollinator foraging in free-flying bees (Dyer et al., 2006; Heran, 1952; Whitney et al., 2008). This is perhaps due to pollinators being unable to attend to smaller differences when foraging and visiting flowers in succession and therefore not experiencing temperature differences between flowers simultaneously.

Increased absolute floral temperature, that is elevated floral surface temperature relative to the environment, and increased within flower temperature contrast, that is, greater differences between the hottest and coldest parts of the flower surface or a more ‘contrasting temperature pattern’, appear to be preferred by bees and lead to higher learning speed and accuracy (Dyer et al., 2006; Hammer et al., 2009; Harrap et al., 2017, 2020b; Whitney et al., 2008). Free-moving bees have shown behavioural responses to differences in floral surface temperature of as little as 2°C in contexts similar to flower foraging (Dyer et al., 2006; Heran, 1952). Thus, floral temperature can influence the foraging success of naïve and experienced pollinators in different ways (Raine and Chittka, 2007, 2008). In turn, this is likely to affect how the insect moves or interacts with the flower (Angioy et al., 2004; Harrap et al., 2020b), and influence pollen deposition and removal (Ashman et al., 2004; Schiestl and Johnson, 2013). Consequently, floral temperature has important ecological consequences, both positive and negative, on the fitness of both plants and the organisms they interact with. However, the extent to which floral temperatures vary with weather conditions and how such variation could impact pollinator behaviour, is not well understood.

Little is known about how weather variables influence floral temperature, other than light conditions from solar radiation (Dietrich and Körner, 2014; Herrera, 1995; Kovac and Stabentheiner, 2011; Rejšková et al., 2010; Rougerie-Durocher et al., 2020; Whitney et al., 2011). During the daytime most flowers maintain elevated floral temperatures above that of the environment to varying extents (reviewed by van der Kooi et al., 2019). Most flower species are non-thermogenic (Seymour and Schultze-Motel, 1997; Seymour et al., 2003a; Seymour and Matthews, 2006; Lamprecht and Seymour, 2010), therefore flowers will mainly warm ‘passively’ with the environment as a result of external environmental influences (*i.e.* weather conditions). Increased sun exposure and increased illumination levels are known to raise floral temperatures. Nevertheless, floral temperatures are affected by transpiration and evaporative water loss (Dakhiya and Green, 2019; Harrap and Rands, 2021a; Patiño and Grace, 2002) which under various weather scenarios could cool the flower. Therefore, environmental conditions influencing these processes, such as environmental temperature, humidity and atmospheric pressure, could indirectly influence floral temperatures by influencing transpiration and evaporation from plants (Gates, 1968). Furthermore, environmental humidity and pressure can have complicated influences on how heat travels through the environment (Minkina and Klecha, 2016; Polezhaev, 2011; Urone and Hinrichs, 2012; Usamentiaga et al., 2014), potentially moderating how flowers warm or cool. Finally, wind speed will also affect heat transfer (Polezhaev, 2011; Urone and Hinrichs, 2012). These environmental variables have not yet been investigated in relation to floral temperature. Here we approach this knowledge gap by measuring how much these variables contribute (if at all) to the variation in observed floral temperatures.

Floral traits may influence the extent to which floral temperature is dependent on weather conditions; potentially explaining differences in floral temperatures and temperature patterns between species for a given set of conditions (as observed by Dietrich and Körner, 2014; Harrap et al., 2017; Rejšková et al., 2010; and Shrestha et al., 2018). Traits that influence the amount of radiation energy intercepted and reflected may play a role in promoting generation of floral temperature. Such traits may include pigmentation (Rejšková et al., 2010; Sapir et al., 2006; Whitney et al., 2011), structural properties (Whitney et al., 2011) including gloss (Whitney et al., 2012), flower shape (Lamprecht et al., 2007, 2006), flower orientation, and solar tracking capacity (Atamian et al., 2016; Totland, 1996; Zhang et al., 2010). Other floral traits may help flowers retain heat or protect it from other cooling influences such as larger flower mass (Dietrich and Körner, 2014), compactness of floral structure (Rands and Harrap, 2021), surface texture (Whitney et al., 2011) and pubescence (Miller, 1986). Such traits can influence surface area to volume ratios and formation of boundary layers and trapped air, thus influencing heat retention and release (Urone and Hinrichs, 2012). Differences in these traits across the flower surface are likely to play an important role in the generation of contrasting temperature patterns (Atamian et al., 2016; Harrap et al., 2017; Lamprecht et al., 2006; Rejšková et al., 2010). Understanding the influence of different floral traits on generation of floral temperature and temperature patterns will further our understanding of how floral temperatures are generated and also identify traits that may be subject to selection mediated by floral temperature’s influences on plant fitness.

Weather-dependent variation in the floral temperature presented by flowers may mean that floral temperature has intermittent or changing ecological impacts. Therefore, understanding how floral temperature varies with weather conditions, the extent of this variation, and the relative importance of different weather variables, is critically important to understanding how often flower temperature has ecological consequences in natural systems. Of particular importance is how floral temperature changes while individual plants interact with their pollinators. This is due to the complex influences that temperature can have on pollinator responses to flowers (van der Kooi et al., 2019). Changing floral temperatures may influence pollinator flower learning, preferences, and handling as well as how rewarding flowers are perceived to be.

Previous work often only measures floral temperature at a single point on the flower, typically on the reproductive structures. How the temperature at different points on the flower surface, temperature patterns varies with weather conditions has not been studied in much detail (but see, Rejšková et al., 2010; and Dietrich and Körner, 2014). Similarly, the effects of variations in conditions such as humidity, pressure and wind on petal temperature are infrequently investigated together (although Rougerie-Durocher et al., 2020 demonstrate that this can be done, in a study measuring apple *Malus pumila* pistils across variable conditions). This may be particularly important for understanding the temperatures that plants show. In this study we use thermal imaging techniques to monitor how absolute floral temperature and temperature patterns of four plant species change with weather conditions during the periods of the day that pollinators are active. We find a strong influence of solar irradiation, as expected, but describe and assess also the effects of other variables.

## Methods

### Study site and species

The study was conducted at the National Botanic Garden of Wales, Carmarthen, UK (51.84N, 4.14W). The flowers of four plant species were monitored, comprising of two wild type species and a pair of closely related horticultural varieties (henceforth ‘species’ collectively).

Rock roses, *Cistus* ‘snow white’ and *Cistus* ‘snow fire’ are horticultural varieties of the genus *Cistus*, which originates in the Mediterranean region (Papaefthimiou et al., 2014). Both varieties produce flowers typical of the white-flowered *Cistus* group (Guzmán and Vargas, 2005; Guzmán et al., 2011) and are visited by bees and flies (typical of *Cistus*, see Manetas and Petropoulou, 2000; Bosch, 2008; Steen and Orvedal Aase, 2011). The two varieties differ in petal base pigmentation but are similar in other regards. While both varieties have white petals with yellow reproductive structures, *Cistus ‘*snow fire’ has dark red-black triangular patches at the base of petals, which are absent in ‘snow white’ (figure 1). Such patches, as well as the absence of such patches, are common across *Cistus* (Guzmán and Vargas, 2005; Guzmán et al., 2011). All *Cistus* flowers were sampled within a *c*. 224m² (approx. 37m by 77m) planted flower bed.

**Figure 1:**
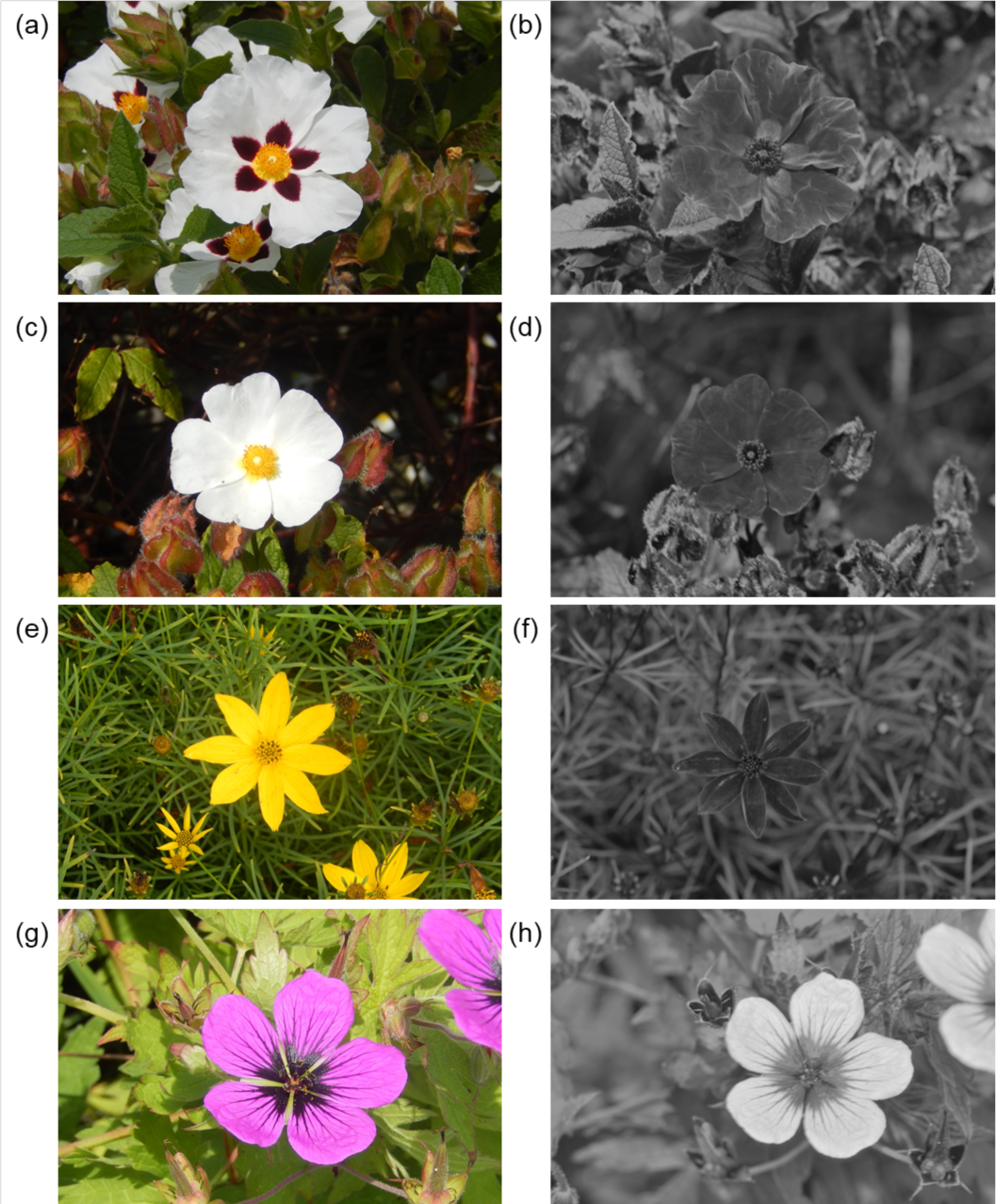
The four species surveyed within the study viewed under human visible (a, c, e, g) light and UV light (b, d, f, h): (a) and (b) *Cistus* ‘snow fire’; (c) and (d) *Cistus* ‘snow white’; (e) and (f) *Coreopsis verticillata*; (g) and (h) *Geranium psilostemon*. UV images were collected using a Nikon D90 with all internal light filter components removed, and a UV permeable lens (UV-MACRO-APO 108012, Coastal Optics) with a filter transmissive to only light of wavelengths 320-380nm (U-Venus-Filter, Baader) fitted externally to the camera. UV images are captured under daylight conditions with approximately a 6 second exposure, to compensate for the relatively low UV light illumination of daylight conditions. UV images are recoloured to black and white for ease of interpretation (the camera as described assigns a colour, normally red, to the UV signals inappropriately, thus only the brightness in these images is appropriate). White areas indicate areas that are UV reflective.

Whorled tickseed, *Coreopsis verticillata* L., is an herbaceous perennial, native to North America but widely cultivated in the UK. Its produces composite inflorescences characteristic of the Asteraceae family (Harris, 1999) with yellow disc florets and yellow petalled ray florets surrounding (figure 1e and f). This composite inflorescence acts as a floral display comparable to a flower in other groups. Like most Asteraceae (Mani and Saravanan, 1999), *Coreopsis* are visited by a range of visitors including bees and flies (Braman et al., 2022). All *C. verticillata* flowers were sampled within a *c*. 2361m² (approx. 12m by 201m) planted flower bed.

The Armenian Geranium, *Geranium psilostemon*, is an herbaceous perennial native to Armenia, Turkey Azerbaijan, and the Russian Federation. It is cultivated and persistently naturalised throughout the UK (Hitchmough and Woudstra, 1999; Stace, 2010). *G. psilostemon* produces flowers with predominantly purple-magenta petals, which also reflect bee-visible UV light, with black petal bases and veins throughout (figure 1g and h). These flowers are visited mainly by bees and flies. The black petal bases and veined areas are glossier due to changes in surface structure between these regions typical of the *Geranium* genus (Whitney et al., 2011, 2012; Papiorek et al., 2014). All *G. psilostemon* flowers sampled grew within a *c.* 10373m² (approx. 82m by 121m) wildflower meadow.

Flowers of each species were monitored during three distinct sampling periods in 2016 and 2017. During these sampling periods floral temperature of particular study species, the ‘focal species’ of that sampling period, was monitored repeatedly during daytime hours across several days. Focal species in each sampling period were either: a) *Cistus* ‘snow fire’ and *Cistus* ‘snow white’; b) *G. psilostemon*; or c) *C. verticillata*. *Cistus* ‘snow fire’ and *Cistus* ‘snow white’ were sampled on the same days, with the exception of one day (05/07/2016) where weather conditions only allowed sampling of *Cistus* ‘snow fire’ (see below for discussion of how weather conditions affected sampling efforts). A summary of sampling periods and numbers of observations are given in table 1, sampling periods were chosen to broadly cover the height of the species’ flowering at the study site.

**Table 1:** A summary of the sampling periods and replication of each flower species surveyed. The start and end dates of each flower species sampling periods in each year are given and the number of days during which sampling took place (weather conditions may prevent continuous sampling every day within the sampling period). Additionally, the overall number of flowers sampled as well as the mean replication on each individual flower is given.

| Species | Year | Start<br>(DD/MM/YY) | End<br>(DD/MM/YY) | Number of<br>sampling<br>days | Number of<br>individual<br>flowers<br>sampled | Replication per<br>flower<br>(mean ± S.D) |
| --- | --- | --- | --- | --- | --- | --- |
| *Cistus*<br>'snow fire' | 2016 | 23/06/16 | 06/07/16 | 11 | 414 | 2.56 ± 1.53 |
|  | 2017 | 20/06/17 | 16/08/17 | 13 |  |  |
| *Cistus*<br>'snow white' | 2016 | 23/06/16 | 06/07/16 | 10 | 357 | 2.59 ± 1.51 |
|  | 2017 | 20/06/17 | 16/08/17 | 13 |  |  |
| *Coreopsis*<br>*verticillata* | 2016 | 10/08/16 | 17/08/16 | 7 | 154 | 5.42 ± 3.60 |
|  | 2017 | 22/07/17 | 30/08/17 | 12 |  |  |
| *Geranium*<br>*psilostemon* | 2016 | 11/07/16 | 31/07/16 | 13 | 238 | 3.79 ± 2.74 |
|  | 2017 | 08/07/17 | 18/07/17 | 7 |  |  |

### Floral monitoring

Floral temperature monitoring only took place between 09:00 and 19:00 under conditions where insect floral visitors were seen foraging on focal species or other plants within the same locations. If conditions deteriorated to a point where there were no active insect visitors present, monitoring would cease. Therefore, floral temperatures were monitored within conditions relevant to pollinator interaction.

Similarly, temperature monitoring was only conducted when conditions were suitable for thermography. If it began to rain or wind speeds increased to a point that flowers would not remain still for long enough for image capture (*c*. 2-3 seconds) monitoring would cease. Normally these conditions coincided with a stop to pollinator activity. If conditions stopped monitoring in these ways, monitoring would resume once conditions improved. This could be at a later point in the same sampling day or the next suitable day. These considerations of conditions also had to be met before the beginning of sampling on the first day of the sampling period.

Sampling commenced with selection of individual flowers of the focal species to monitor. Upon selection, flowers underwent thermographic sampling (capture of thermal images and associated measurements) using the procedure described in the following section. Flowers were not selected if they were visibly damaged, diseased or had standing water visible on the flowers. Flowers could be selected if they had an atypical number of petals (for example, six-petalled *Cistus* flowers, where five-petalled flowers are typical) if this clearly reflected natural variation in flowers and not damage. Flowers had to be accessible with thermal imaging equipment without having to climb or stand upon other plants in managed plots or, when sampling *G. psilostemon* within wildflower meadows. Where flowers were clustered in such a way that picking out individuals was difficult, it was more convenient to sample all flowers in the cluster (effectively adding extra flowers to the subset) and thus avoid potentially missing those sampled previously due to misidentification. Flowers were not directly handled or moved to allow imaging, as this would influence temperature. Therefore, most of the flower’s upper surface had to be viewable at a right angle to the flower’s span. Additionally, any flowers selected that were being monitored at one time came from across at least two locations at least 2 m apart within their sampling area. Selection was otherwise haphazard, from those flowers available. We continued to select and thermograph flowers until a subset of approximately fifteen (in 2016) or ten (in 2017) individual flowers of each the focal species had been sampled (above caveats for clusters notwithstanding). As both *Cistus* varieties were sampled simultaneously (see table 1), here 15 (in 2016) and 10 (in 2017) flowers of both varieties, so a maximum of 30 or 20 flowers in total, were monitored at a given time, dependent upon the availability of flowers.

Once this subset of flowers had been thermographed, we repeated thermographic measurements of this subset throughout the day replacing flowers as required. When repeat measurements were conducted, where possible we returned to the same flowers selected previously and thermographed them. If previously selected flowers had visible standing water on them (for example from condensation or rain occurring between repeats), they were not thermographed on that repeat, because the thermal camera would measure the temperature of the water surface on the flower as opposed to that of the floral tissue. If standing water was no longer visible at a subsequent repeat measurement, thermography of that flower was resumed. Similarly, if a *C. verticillata* flower had closed, it was not thermographed but monitoring would resume if it opened on subsequent repeats. If, by the time of repeat thermographic measurements, a selected flower had: become damaged, diseased or wilted; begun to develop fruits; abscised petals; or simply could not be found, a new flower was selected as described above to replace it. If no open flowers were available at that time, no replacement occurred. In the case of *G. psilostemon* and *C. verticillata*, selected flowers were sampled across several days, with previously selected flowers being returned to where possible and new flowers being selected as required. In the case of *Cistus* where flowers last less than a day, replacement was carried out throughout the day, but by the end of a sampling day we would reach a point where there were no available replacement flowers. Thus, while monitoring *Cistus*, at the beginning of each sampling day a whole new subset would have to selected.

The public location of the plants meant that it was not possible to label individual plants, but it was easy for the thermographer (see author contributions) to memorise their locations for repeated measurements using positional cues. Furthermore, the identity of flowers was confirmed by comparing captured thermal images with the human-visible colour image that was taken at the same time by the thermal camera (see below). This meant that the likelihood for errors in individual flower identification was low.

### Floral thermograph procedure

Thermography techniques (Usamentiaga et al., 2014; Tattersall, 2016; Vollmer and Möllmann, 2017), were used to monitor floral temperature. The protocols used are in line with established best practices (Harrap et al., 2018) and have been used previously to evaluate and monitor floral temperature traits (Lamprecht et al., 2006, 2007; Lamprecht and Seymour, 2010; Rejšková et al., 2010; Dietrich and Körner, 2014; Faye et al., 2016; Harrap et al., 2017; Dakhiya and Green, 2019; Byerlay et al., 2020; Rougerie-Durocher et al., 2020; Harrap and Rands, 2021b), All thermal imaging was conducted by a qualified thermographer (see author contributions) with a FLIR E60bx thermal camera (FLIR systems, Inc., Wilsonville, USA). The camera can also record human-visible colour images together with each thermal image and assigns a date and time taken to each image.

Thermal images were taken of flowers, or inflorescences in the case of *C. verticillata*, facing their upper surface at a right angle to the flowers span (figure 2). Thermographs were taken at approximately 50cm distance, which ensures all visible parts of the flower’s upper surface could be viewed in a single image. The thermographer avoided standing in positions that cast shade on a flower as much as possible.

**Figure 2:**
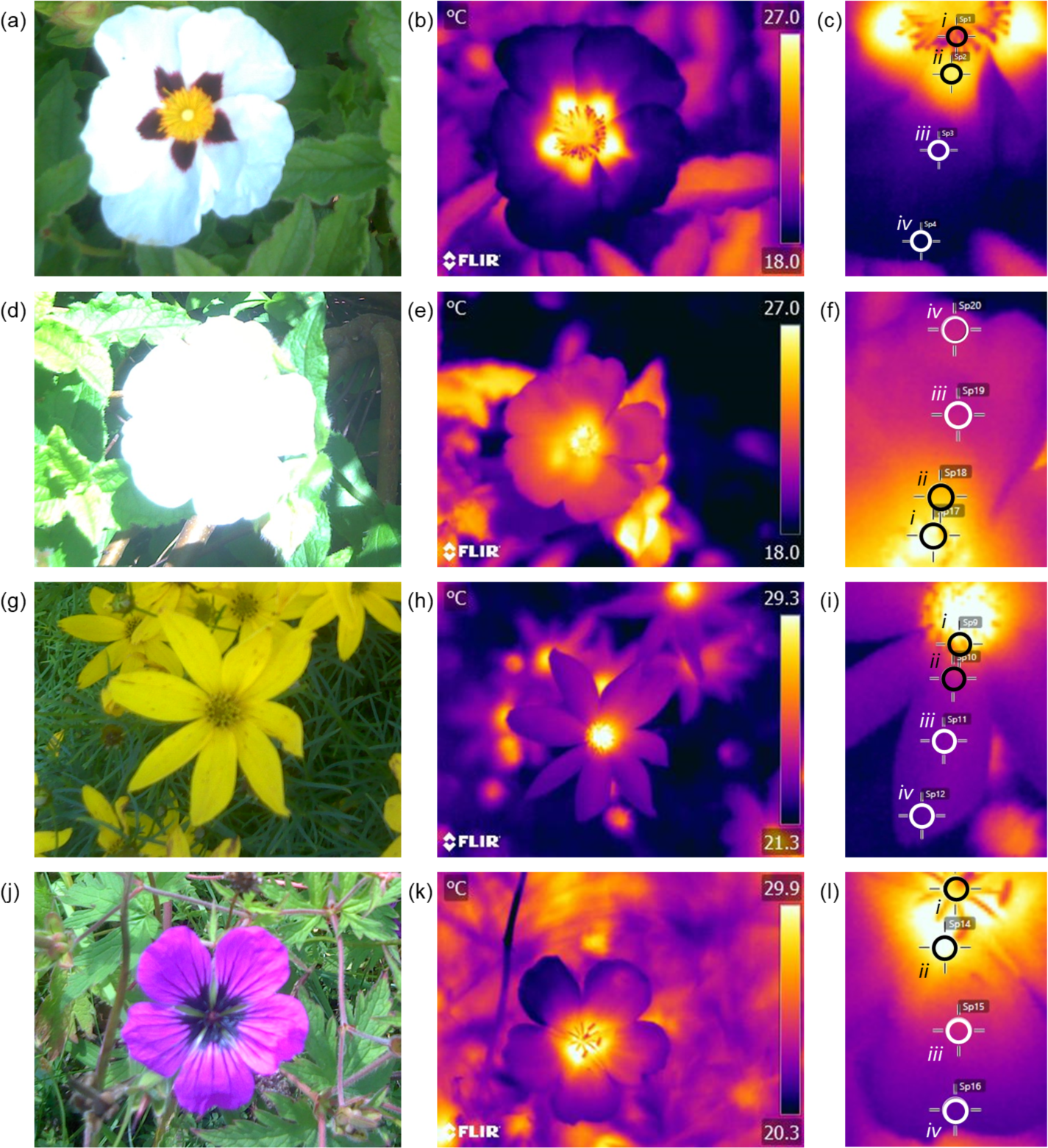
Example thermal images of (a, b, c) *Cistus* ‘snow fire’, (d, e, f) *Cistus* ‘snow white’, (g, h, i) *Coreopsis verticillata* and (j, k, l) *Geranium psilostemon*. Human colour image captured by the thermal camera (a, d, g, j) are provided for reference (not this camera cannot have focus or exposure adjusted). Thermal images are provided as captured (b, e, h, k) for each species. Temperature is given by the colour scale to the right of thermal images in degrees Celsius. Lastly the floral temperature measurement points corresponding to one petal are given as a screen capture from *FLIR tools* (c, f, i, l). *FLIR tools* measurement superimposed with black or white rings that contrast with thermal image coloration for heightened clarity. Numerals on screen capture indicate position of thermal measurement on flower: *i.* Reproductive structures, *ii.* Petal base, *iii.* Petal middle, and *iv.* Petal tip. Specific thermograph details and environmental conditions of thermographs are discussed in Supplementary material 1.

The procedure was as follows for each flower and took approximately 4 minutes. First, a thermal image was recorded, from which afterwards flower temperature measurements were later taken. Accurate thermographic temperature measurements require well-focussed thermal images to be captured (Vollmer and Möllmann, 2017; Harrap et al., 2018; Usamentiaga et al., 2014). Multiple thermal images were often captured in a short sequence to ensure a well-focussed image had been taken (normally two or three attempts). In order to later extract the reflected temperature (a key thermography parameter) for each thermograph, a thermal image was recorded of a reflective object (a tin foil multidirectional mirror, *c.* 15 × 7 cm) that was placed next or on the flower (this image is referred to as the ‘reflected temperature image’ below). At the same time, illumination (lux) was measured using a lux meter (*KKmoon* HS1010, Shenzhen, China) held next to the flower matching the orientation of the flower’s upper surface. The mirror had text identifying the flower written on it and the lux value to facilitate differentiation between samples and matching Lux values with thermographs.

When flowers selected for monitoring were very close to each other (approximately less than 10 cm) and of similar orientation, such that the multidirectional mirror would have to be placed in the same position for each flower, each of those flowers was imaged, and a shared reflected temperature measurement was used. These flowers also normally shared lux measurements if the lux meter likewise was placed in the same position for both flowers. Note that, as flowers can change their orientation with time (due to growth or wind, *etc*.) flowers that shared lux or reflected temperature measurements at one sampling event did not necessarily share the measurements in other sampling events conducted earlier or later.

### Image processing

Following collection, images were sorted and matched by individual flowers across the sampling period, using colour images and labels on the corresponding reflective temperature images. If a flower could not be reliably identified, it was assumed to be a new flower. Where multiple images of the same flower were taken at the same sampling event (to ensure a focused thermal image, see above) only one thermograph was selected for each sampling of each flower. A few images were discarded due to poor image quality, in terms of focus and image capture of the flower’s surface.

Floral temperature measurements from thermographs were made using *FLIR tools* (FLIR systems INC, 2015). For floral thermographic temperature measurements, emissivity was set to 0.98. This value is suitable for floral emissivity across locations and species (Harrap and Rands, 2021b). Reflected temperature for each measurement incident was obtained from the corresponding reflected temperature image of the multidirectional mirror, taken after each flower thermograph (see above). Reflected temperature was measured by taking the average apparent temperature of the mirror in this thermal image (obtained by setting emissivity and distance parameters in the mirror thermograph to 1 and 0 respectively). Other thermography parameters for floral temperature measurements were set as follows: distance 0.5m, environmental temperature 25°C, relative humidity 50%. These ‘other’ thermography parameters account for transmissivity of the air between the camera and target and radiation from that air. Transmissivity is normally near 100%, and radiation of the air is small unless environmental temperature or humidity are at extreme values, or images are captured over long distances (Minkina and Klecha, 2016). Although environmental temperature and humidity changed between measurement incidents, because images were captured over very short distances the effects of using fixed values for these ‘other’ parameters on floral temperature measurements would be minor (Usamentiaga et al., 2014; Vollmer and Möllmann, 2017; Harrap et al., 2018).

On each selected flower thermograph, temperature measurements were taken using the point measurement function in *FLIR tools*. Measurement points were placed manually. Point measurement functions on thermal cameras have a minimum area they measure and maintain accuracy indicated automatically in *FLIR tools*. Consequently, points were placed such that the entirety of the measurement point was on the relevant structure (figure 2). Points were placed so that they had minimal overlap with each other (this rarely occurred outside of reproductive structures measurements). Measurement point placement was determined on what is visible in the image, thus if part of the flower was obscured, for example by foliage covering the very tip of the petal, or by other parts of the flower itself (due to off-centre image alignment), point placement was adjusted to what remained visible in the image. If in a thermograph a petal was completely obscured in this way, measurements were not taken from that petal.

Floral temperature was measured at four positions across the flower (figure 2): on the reproductive structures (pistils, stamens, disc florets) and at the base, middle and tip of the petals. “Reproductive structure” measurements were taken at a point corresponding to each petal of the flower in each thermograph, taken at the point visible on the image nearest each petal but remaining entirely on the reproductive structures. The exception to this was the reproductive structure measurements on *G. psilostemon*. The small size of *G. psilostemon* reproductive structures relative to petals made it difficult to place multiple measurements points on them without these points overlapping completely. Thus, a single reproductive structure measurement was taken on each *G. psilostemon* thermograph at the centre of the reproductive structures visible in the image (figure 2).

Petal measurements were taken on each petal of the flower in each thermograph, or on the ray floret petals in *C. verticillata*. The petal base and tip measurements were placed at the point visible in the thermal image as close to the base and tip that point placement would allow. The petal middle measurement was taken at a point halfway between these points. With consideration for minimum size for point placement, this effectively meant the ‘petal base’ measurement corresponded to a position within the first 25% of the petal’s length (going from base to tip), the ‘petal middle’ a point approximately 50% along the petal’s length, and the petal tip measurement, a position within the last 25% of the petal’s length. This meant the darker locations of *Cistus* ‘snow fire’ and *Geranium psilostemon* petals were both the location of petal base measurements. Where possible, the petal measurements were placed along the central line of the petal but would be offset from this centre line if that position was obscured.

### Weather conditions

During floral monitoring, the illumination measurements (lux/100) at each thermography incident were obtained (described above). Hourly weather data at the time of each thermographic measurement was obtained from the Pembrey Sands weather station (51.71N, 4.37W) 21.4km away from the Carmarthen study site, *via* the UK Met office. From this, the following weather measurements for each hour of the day during sampling were obtained: hourly temperature (°C); hourly mean wind direction (degrees); hourly mean windspeed (kn); hourly total cloud cover (oktas); hourly relative humidity (%) and hourly pressure at mean sea level (hPa). The Pembrey Sands weather station provided consistent monitoring for each of these variables across all hours where thermography took place, with the exception of hourly total cloud cover readings for a single hour (09:00 on 16/7/18) where the value was missing, during which *G. psilostemon* was sampled. Given the hourly total cloud cover readings for the hour before and after this both read as 10 oktas, we assumed cloud cover did not change over this time period and substituted 10 oktas for the missing cloud cover reading of this hour. For the purposes of assigning a time to the thermography incident, the time of the reflected temperature image capture was used (as recorded by the thermal camera). Time of day itself was included in our analyses as ‘the hour of the day of each thermography incident’ as an additional variable.

Hourly mean wind direction was considered in our analyses of correlation of conditions (see supplementary materials). However, the nature of conditions that correspond with wind direction will vary greatly from location to location and mean wind direction had significant relationships with all other weather variables (see supplementary materials). Consequently, it was deemed more informative to not include wind direction in our further analysis, whose effects are likely region-specific or reflect changes in other conditions with wind direction. Additionally, this exclusion reduced the complexity of statistical analyses. This left us with 6 weather variables and time of day (7 total).

### Statistical analysis

#### Weather variables influence on floral temperature

Correlation between all six measured weather variables and time of day was assessed (Supplementary information 2). As expected with weather variables, there were statistically significant covariation between many of the weather variables but these correlations were of weak to moderate strength (−0.4 < *r* < 0.4, see supp figure S1). The methods used for analysis of weather influences on floral temperature (model comparisons based on AIC) are robust to moderate covariation between variables (where −0.5 < *r* < 0.5, according to Freckleton, 2011). Consequently, the covariation was deemed unlikely to interfere with model selection. Whether study species differed in the conditions experienced was assessed (Supplementary information 3). Frequently the two *Cistus* varieties experienced (statistically) similar conditions to each other, while in some variables *G. psilostomon* and *C. verticillata* experienced similar conditions, although these groupings still differed from each other. Although species differed statistically in the conditions they experienced, these differences were not large and species experienced conditions across a comparable range for each weather variable (Supplementary information 3). Thus, it was considered unlikely that differences in conditions experienced by species would alter the effects on floral temperatures identified by AIC models.

The effects of each of the six included weather variables, and time of day, on floral temperature for each of four positions on the flower of each species was assessed independently using AIC model simplification techniques (Richards, 2008). This analysis was conducted in *R* 3.6.3 (R Core Team, 2020), utilizing the package *lme4* 1.1.-25 (Bates et al., 2015). This involved paired AIC comparisons between a standing best model and a simpler model, where simpler models were constructed by removing parameters from the standing best model (by forcing those parameters to be zero). If removal of parameters resulted in a sufficient increase in AIC, based on Richards (2008), the standing best (more complex) model would remain the best for the next comparison. If otherwise, the simpler model would become the standing best model for the next comparison. Initially, a model allowing flower position and each weather variable to influence floral temperature and temperature patterns was fit to the data of each species, the ‘full model’. This full model allowed a quadratic effect of time of day (expressed as decimal hour of the day after 09:00, thus 09:00 = 0, 10:00 = 1, *etc*.), a logarithmic effect of illumination, and linear effects of all other weather variables on floral temperature. This full model did not allow weather variables and time to have interacting effects on floral temperature but did allow position on the flower to have interacting effects with weather variables. These interactions between variables and position on the flower describe the variables influence on the flower’s temperature pattern (*i.e.* how different positions on the flower warm or cool). Flower position could also influence floral temperature independently of weather conditions (influencing the model’s intercept). A full description of the full model and simpler models is given in supplementary information 4.

The sequence of AIC comparisons testing the effect of each weather variable on floral temperatures was conducted in the same order in each flower species: wind speed, atmospheric pressure, relative humidity, cloud cover, environmental temperature and illumination. For each weather variable, the effect of removing the parameters that allow an interaction between that variable and flower position was tested first. This first comparison allows assessment of whether the variable influences temperature pattern contrast and structure. Following this, the effect of the variable on absolute floral temperature (that is floral surface temperature independent of position) was tested. This was done by removing all parameters that allow floral temperature effects in response to that that weather variable within the standing best model. This meant, if the position-dependent effects had been removed following the previous comparison (based on the AIC comparison), removing the remaining position-independent effect of that variable to make the simpler model for this comparison. If position-dependent effects of that variable were maintained in the standing best model after the previous comparison, these position-dependent effects were removed along with the position-independent effect. Once these comparisons had been conducted for each weather variable, the effect of time of day was tested in the same way (position-dependent effects then absolute effects). Following this, the weather-independent effects of flower position (positions’ intercept effects) were tested.

Testing absolute effects when position-dependent effects are known to be important (*i.e*. already demonstrated to be retained in the best model) could be considered a redundant test in terms of understanding what variables should be included in the final model, and therefore what variables influence floral temperature for that species. That is, if we know floral temperature patterns are affected by the variable in question, absolute floral temperature is affected by it. However, the loss of information (represented by ΔAIC) between standing best and simpler models in this comparison still provides information on the impact and importance of the weather variable on absolute floral temperature. Furthermore, in these instances, the difference in information loss between comparisons of position-dependent and absolute effects provides an indication of the position-independent impact of the variable on floral temperature (a value comparable to the other instances where position-dependent effects are already removed).

#### Flower positions influence on temperature’s weather variable response

The standing best models for each species after the last of these comparisons was then used to evaluate how much flower positions differ in their responses to weather conditions and thus floral temperature. Alternative versions of the standing best model (maintaining the same model structure and effects) with alternative groupings of the flower position factors would be fit to the data of each species. These alternative groupings treated positions grouped together as the same, effectively as a repeat measurement of a single combined flower position. These alternative groupings included all possible grouping combinations of the four positions. This included a model where all positions are treated as the same (where all position effects were simply removed) and a model where all positions differed, which was the standing best model at the end of the comparisons to determine weather effects. These models are described in more detail in Supplementary information 4. These ‘alternative position models’ were compared with AIC. The best fitting model at the end of this AIC comparison was considered the best model of floral temperature for each species.

## Results

### Weather variables influence on floral temperature

In all species, floral temperature and contrasts in temperature between different positions on the flower were influenced by multiple weather variables. The results of the model selection process for the floral temperature responses of each flower species to weather conditions are summarised in tables S4 to S9 (see Supplementary Material). Figures 3-6 show the effects of each weather variable on the floral temperatures, according to our final best models, for *Cistus* ‘snow fire’*, Cistus* ‘snow white’*, C. verticillata* and *G. psilostemon* respectively. The results shown are based on the model effects. Information described by the variable is used to evaluate the importance of each variable; this is indicated by model comparisons, ΔAIC, when the parameter is removed from models, see tables S4 to S7 in the Supplementary materials.

**Figure 3:**
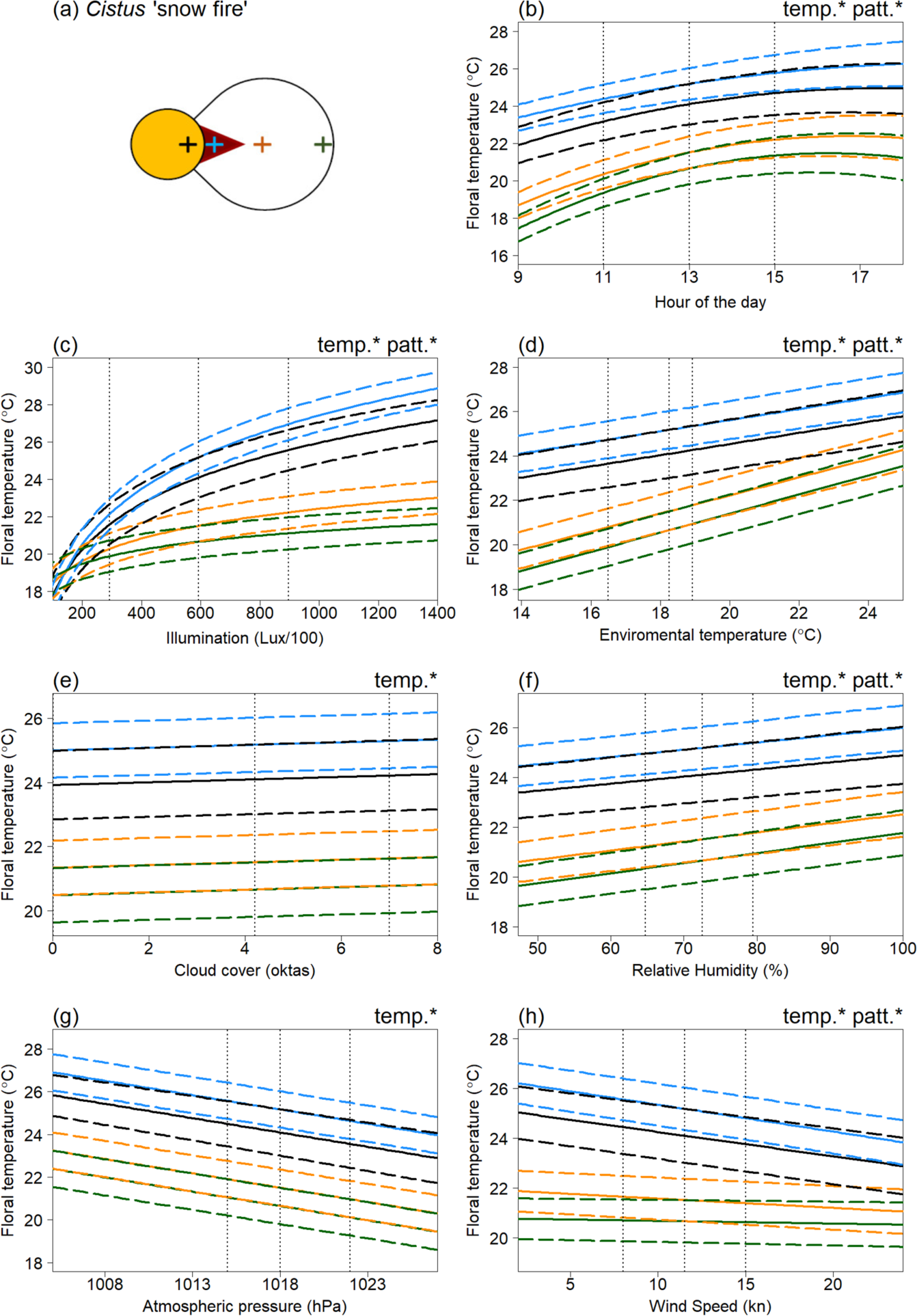
The effects of weather conditions on the floral temperature of *Cistus* ‘snow fire’ according to our best model of floral temperature. The influence of changes in (b) hour of the day, (c) illumination at time of imaging, (d) hourly environmental temperature (e) hourly cloud cover (f) hourly relative humidity (g) hourly atmospheric pressure, and (h) hourly wind speed, is shown for mean conditions (from across all the sampling period) for all other weather variables and during the 13^th^ hour of the day (13:00, true mean hour across sampling was 12:46). Line colour indicates location on the flower: ‘black’, the reproductive structures; ‘blue’ the petal base; ‘orange’ the petal middle; ‘green’ the petal tip. These locations are indicated by crosses of the same colour on the diagram of the species petal and reproductive structures in panel (a). The mean temperature of each flower location as described by the best model is indicated by bold solid lines, and ±0.5 S.E.M by long-dash lines. Vertical dotted lines indicate (from left to right) the first quartile, mean and third quartile conditions for each weather variable across the whole of the sampling (hour 13 is taken for mean hour of the day, see above). Note first quartile cloud cover is 0 oktas, this line is offset of this position slightly to be made visible. Conditions at the middle, mean, vertical line are the same across all panels. ‘temp.*’ and ‘patt.*’ in panel corners indicate the corresponding variable influences absolute floral temperature (temp*) and the contrast of the temperature pattern (*patt) respectively according to our best fitting model.

**Figure 4:**
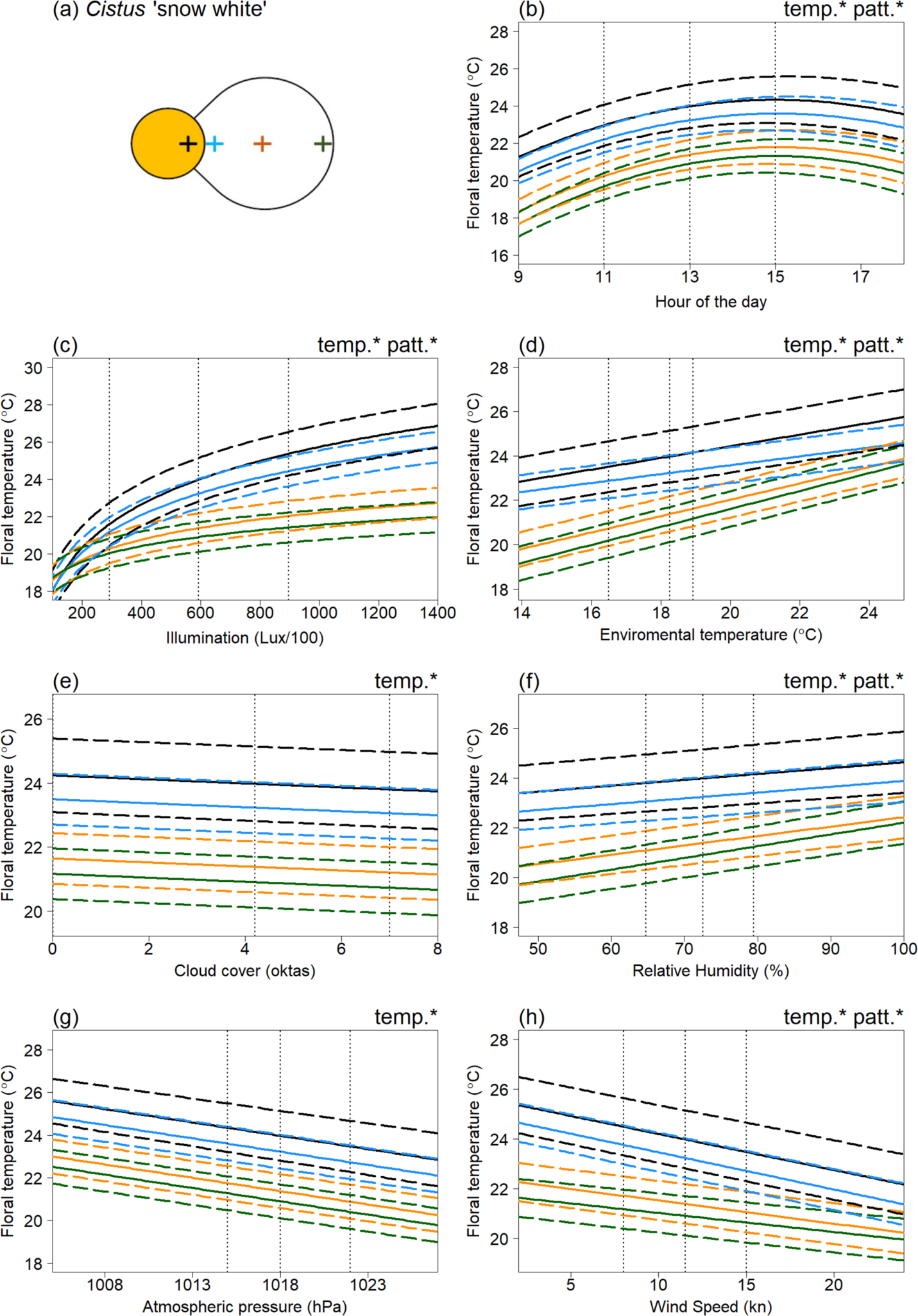
The effects of weather conditions on the floral temperature of *Cistus* ‘snow white’ according to our best model of floral temperature. The influence of changes in (b) hour of the day, (c) illumination at time of imaging, (d) hourly environmental temperature (e) hourly cloud cover (f) hourly relative humidity (g) hourly atmospheric pressure, and (h) hourly wind speed, is shown for mean conditions (from across all the sampling periods) for all other weather variables and during the 13^th^ hour of the day (13:00, true mean hour across sampling was 12:46). Details otherwise as in figure 3.

**Figure 5:**
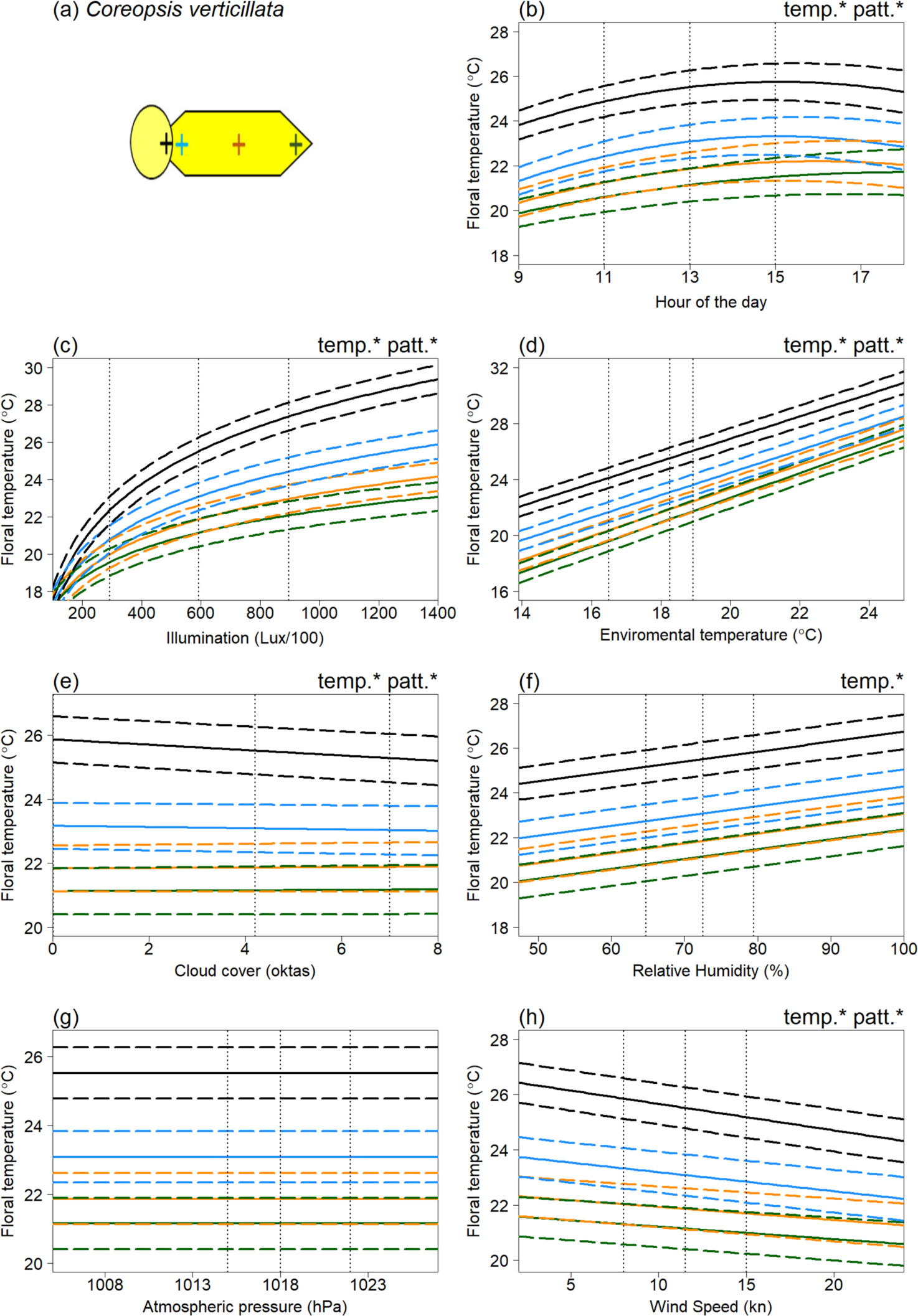
The effects of weather conditions on the floral temperature of *Coreopsis verticillata* according to our best model of floral temperature. The influence of changes in (b) hour of the day, (c) illumination at time of imaging, (d) hourly environmental temperature (e) hourly cloud cover (f) hourly relative humidity (g) hourly atmospheric pressure, and (h) hourly wind speed, is shown for mean conditions (from across all of the sampling periods) for all other weather variables and during the 13^th^ hour of the day (13:00, true mean hour across sampling was 12:46). Details otherwise as in figure 3.

**Figure 6:**
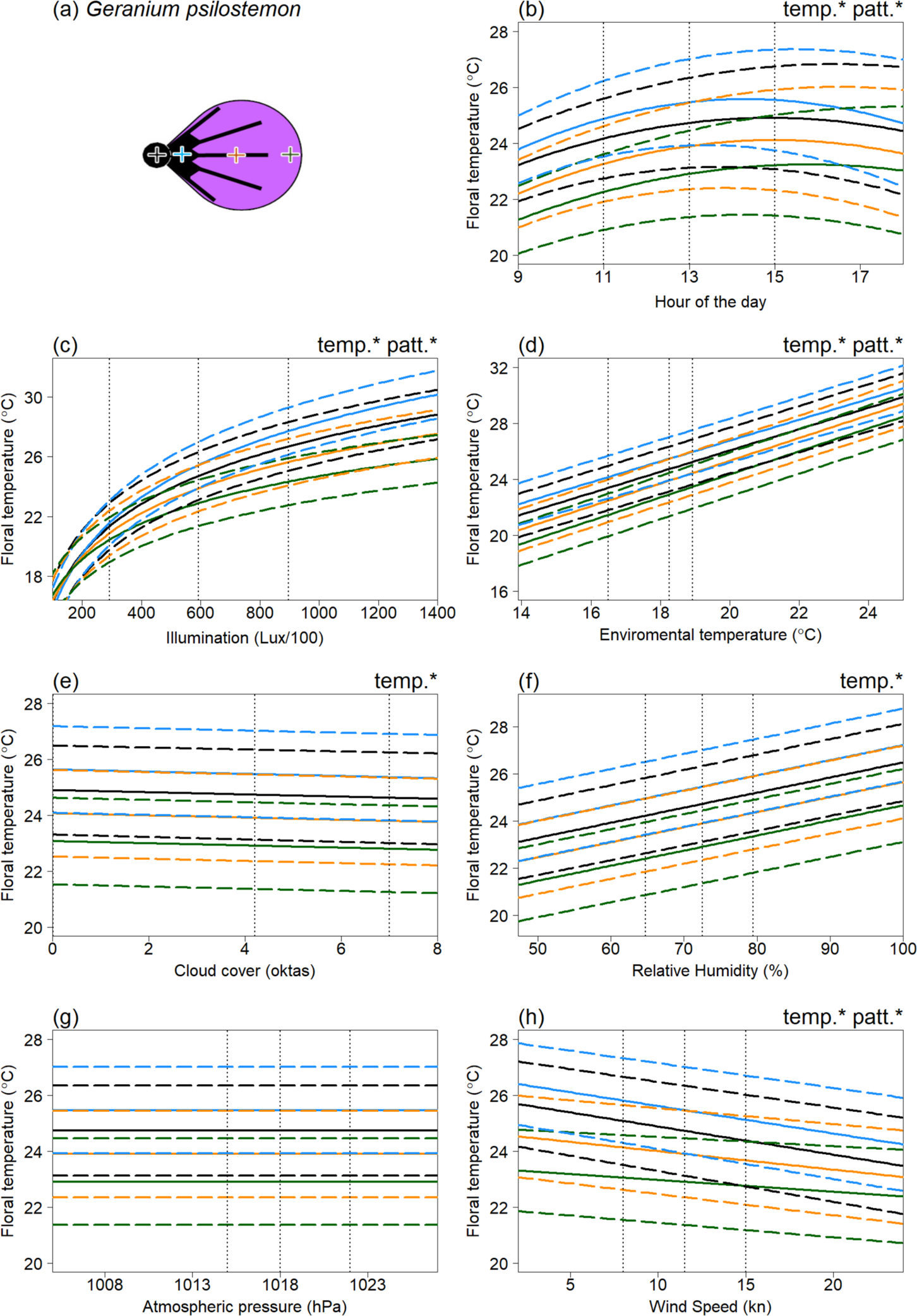
The effects of weather conditions on the floral temperature of *Geranium psilostemon* according to our best model of floral temperature. The influence of changes in (b) hour of the day, (c) illumination at time of imaging, (d) hourly environmental temperature (e) hourly cloud cover (f) hourly relative humidity (g) hourly atmospheric pressure, and (h) hourly wind speed, is shown for mean conditions (from across all of the sampling periods) for all other weather variables and during the 13^th^ hour of the day (13:00, true mean hour across sampling was 12:46. Details otherwise as in figure 3.

Absolute floral temperature (surface temperature independent of flower position) was influenced by all weather variables and time of day in both *Cistus* varieties (figures 3 and 4), and by time of day and weather variables except atmospheric pressure in *C. verticillata* and *G. psilostemon* (figures 5 and 6). Absolute floral temperature of all flower species was most influenced by illumination. In both *Cistus* varieties, time of day had the second most important effect on absolute floral temperature, with environmental temperature being third most important for *Cistus* ‘snow fire’ and wind speed third for *Cistus* ‘snow white’. In *C. verticillata* and *G. psilostemon* environmental temperature had the second greatest effect, while time of day had the third.

Temperature differences between flower positions, that is ‘temperature patterns’ (position-dependent temperature effect), in both *Cistus* varieties was influenced by time of day and all weather variables except atmospheric pressure and cloud cover (figures 3 and 4). *C. verticillata* temperature patterns were influenced time of day, illumination, environmental temperature, cloud cover and wind speed (figure 5). *G. psilostemon* temperature patterns were influenced by time of day, illumination, environmental temperature, and wind speed (figure 6). Temperature patterns of *Cistus* ‘snow fire’*, Cistus* ‘snow white’ and *G. psilostemon* were most influenced by illumination and then by environmental temperature. Wind speed had the third largest effect on temperature patterns in *Cistus* ‘snow white’ and *G. psilostemon*. Time of day had the third most important effect in *Cistus* ‘snow fire’. In *C. verticillata*, illumination, cloud cover and wind speed (in that order) had the most important effects on temperature patterns.

### Position-dependent temperature responses to weather variables

Across all species, models that allowed each position on the flower to differ from all other positions in their responses to weather conditions, time and in their intercept effects (the ‘full model’ with no grouping of different positions) had lower AIC than all other alternative position models, which grouped the different flower positions together. This indicates that each flower position differs in their response to weather conditions, and therefore also the floral temperatures realised for given conditions, in all four species. The full results of the AIC comparisons of all models with alternative position groupings are given for each flower species in table S8 in the Supplementary material.

## Discussion

Under natural conditions, floral temperature and temperature patterns of all species monitored were observed to increase and decrease under various weather conditions. Although species differed in the extent to which they showed elevated floral temperatures and within flower temperature contrasts as a function of different weather variables, the relationships uncovered followed our expectations derived from previous studies looking at the weather variables in isolation (see Dakhiya and Green, 2019; Dietrich and Körner, 2014; Harrap and Rands, 2021a; Herrera, 1995; Kovac and Stabentheiner, 2011; Patiño and Grace, 2002; Rejšková et al., 2010; Rougerie-Durocher et al., 2020; and Whitney et al., 2011). Despite some differences between species, there were clear common trends in temperature-weather variable relationships, specifically the directionality (positive or negative) of relationships between each weather variable and floral temperature and the order of impact of the different variables.

### Illumination and light conditions: critical for elevated floral temperature

Flowers showed increased temperatures with increased illumination, environmental temperature and humidity. The relationships between environmental temperature and humidity and floral temperature appeared to represent ‘passive’ warming of the flower in pace with the environment. Without increased illumination (such as when Lux/100<200), flowers were limited in their ability to heat beyond environmental temperatures or generate contrasting temperature patterns (see figures 3 to 6 and S9), maintaining temperatures comparable to the environment (18.3°C in figures 3c, 4c, 5c and 6c) across the flower. However, floral temperature and the contrast in floral temperature between positions increased with relatively small increases in Lux, resulting in elevated floral temperatures relative to the environment once flowers were exposed to moderately sunny conditions (>300 Lux/100). The results confirm that generation of elevated absolute floral temperatures and contrasting temperature patterns is heavily dependent on illumination in all of the species monitored, although flowers were also warmer with increased environmental temperature, and humidity. This means that changes in light conditions, which will be particularly driven by sun exposure, have large influences on the formation of elevated floral temperatures and temperature pattern contrast. This is in line with previous findings and suggestions for non-thermogenic flowers (e.g. Herrera, 1995; Dietrich and Körner, 2014; Kovac and Stabentheiner, 2011; Rejšková et al., 2010; Rougerie-Durocher et al., 2020; Whitney et al., 2011). The critical influence of illumination, and therefore sun and light conditions, can be easily explained; by interception of solar radiation by flowers, with greater illumination, more energy from radiation (light) can be intercepted by flowers leading to elevated floral temperatures.

Flower illumination measurements, unsurprisingly, show a moderate negative correlation with cloud cover (Supplementary figure S1), explaining its negative effects across most species. However, in *Cistus* ‘snow fire’ cloud cover had a very weak positive effect in the final model. The lower importance of cloud cover relative to other variables and its positive effect in *Cistus* ‘snow fire’ could be considered contradictory to the apparent importance and effects of light conditions indicated by floral temperature’s relationship with illumination. These meagre cloud cover effects are likely to be a consequence of illumination measurements better describing the light conditions a flower experiences. Furthermore, hourly cloud cover over an area does not necessarily represent the amount of light reaching flowers as accurately as illumination measurements, that were captured at each flower after temperature measurement. For example, flowers may be in shade, thus experiencing low illumination regardless of cloud cover, or illuminated in a temporary break in the clouds resulting in the reverse.

### Illumination and light conditions: responses moderated by floral pigmentation

In our study the strength of the relationship between floral temperatures and illumination/light appeared to be influenced by pigmentation of flowers, that is the colour of the flower surface, although further work is needed to establish this more firmly. Darker pigmented regions of flowers, both between and within species, warmed up more than other parts of flowers. This is consistent with the effect pigment and colour has on the reflection and absorption of radiation and consequently its capacity to warm at a given illumination, darker pigments reflecting less and absorbing more radiation leading to increased warming and the reverse with lighter pigments (Whitney et al., 2011). The importance of pigmentation is particularly true in terms of the generation of temperature patterns, which often followed patterns of contrasting pigmentation within individual flowers. In the two *Cistus* varieties, where flowers are similar other than petal base pigmentation (figure 1), temperature measurements in the areas of the reproductive structures and of white parts of the petals were comparable between the two varieties. However, the dark petal bases of ‘snow fire’ achieved greater temperatures with greater illumination, that is at higher light levels (compare figures 3C and 4C). This resulted in ‘snow fire’ reaching a greater absolute floral temperature at a given level of illumination. Also, within-flower temperature contrasts were greater when compared to ‘snow white’ at a given level of illumination. Furthermore, the presence of dark pigmentation, and its effect on warming with illumination, altered the arrangement of elevated floral temperature between the two *Cistus* varieties, and so altered the shapes of floral temperature patterns generated under illumination. Temperature patterns of ‘snow white’ followed a gradient from the flower centre, at the reproductive structures, to the flower periphery (a pattern of heating likely the result of flower structure in absence of pigmentation, discussed below), while in ‘snow fire’, the hottest parts of the flower were the dark petal bases, followed by the flower centre (see figure 2, and compare petal base measurements in figure 3C and 4C). Temperature patterns in ‘snow fire’ showed a ring of elevated temperature at the petal base, while other positions followed the same gradient as ‘snow white’.

Similarly, petal pigmentation influences on floral temperature and temperature patterns are seen also in *G. psilostemon*, which is more darkly pigmented than the other species monitored (figure 1), and consequently achieves greater absolute temperatures at a given illumination (figure 6). Furthermore, the black regions of *G. psilostemon* (the reproductive structures and petal bases) show higher temperatures than the more reflective purple-UV regions of its petals, with areas of elevated temperature corresponding with darker regions. The more homogeneous darker pigmentation across the *G. psilostemon* flower could well be the reason for the observed lower within-flower temperature contrasts than flowers such as Cistus ‘snow fire’. This may mean *G. psilostemon,* produces absolute floral temperature cues that contrast well with the environment or cooler conspecific flowers and more efficiently warm visitors. Such absolute floral temperature cues may be more salient to pollinators and preferred by them (Dyer et al., 2006; Hammer et al., 2009; Rands and Whitney, 2008; Whitney et al., 2008). However, *G. psilostemon* temperature pattern cues may be less salient and less informative due to the lower within-flower temperature contrast (Harrap et al., 2017, 2020b). Interestingly, the presence of other petal reflective properties in *G. psilostemon*, petal gloss at the otherwise dark petal base regions, did not appear to have any impact, with darker glossy regions at the petal base still warming rapidly at higher light levels (figure 6C).

### Illumination and light conditions: responses moderated by floral structure

Although pigmentation played a key role in determining the rate of floral warming with illumination, and generating temperature patterns, there remained differences in response to illumination, particularly between different flower positions, that could not be explained by pigmentation differences alone, suggesting there are other influences on floral warming with increased light conditions. We observed that reproductive structures of flowers at the centre of the flower displayed higher temperatures than petals under increased illumination, even when they were not darkly pigmented such as in *Coreopsis* and *Cistus* ‘snow white’. Furthermore, positions further from the flower centre generally showed reduced responses relative to similarly pigmented positions nearer the centre. Temperature patterns of all flowers showed some gradual contrast between similarly coloured regions at the centre to the periphery (figures 2 to 6), but to differing extents.

These remaining responses to illumination possibly reflect the roles of flower shape, structure and geometry on the flower’s capacity to intercept heat for solar radiation and subsequently retain this heat. These factors may explain the elevated warming of the reproductive structures of *Coreopsis* with increased illumination compared to its petals despite similar pigmentation across the flower. This species, as all others across the Asteraceae, have compound inflorescences (Harris, 1999), the reproductive structures measurements corresponding with the disc florets. The disc florets have a tightly compact structure which may facilitate interception of solar radiation and heat retention. We would expect these structural effects on floral temperature under different light conditions to occur across the Asteraceae family, thus this influence of structure on warming under illumination may explain common trends for highly contrasting floral temperature patterns in this family (Rands and Harrap, 2021).

Flower structure may similarly explain the observed gradients in warming from the flower centre to the periphery. It is likely that warming of the enclosed flower centre and areas of dark pigmentation of flowers (discussed above where they are present) transmit heat to other flower positions via conduction. Creating a gradient in temperature, a temperature pattern, from warmer ‘heat source’ areas of the flower to the regions that are cooler, a phenomenon observed previously by Rejšková et al. (2010). This may explain why areas closer to the flower centre, and therefore these ‘heat sources’, show increased temperature with illumination relative to otherwise similar areas towards the flower periphery. Such floral structure and geometry influences are often not considered when assessing floral temperature (van der Kooi et al., 2019). However, the presence of changes in temperature with illumination independent of darker pigmentation, seen here in all species, suggest their importance in understanding formation of elevated floral temperature and particularly contrasting temperature patterns.

### Other weather variables have lesser influences on floral temperature

Aside from illumination other weather variables, namely: environmental temperature and humidity; wind speed; cloud cover; and atmospheric pressure, influenced floral temperature. However, the influence of these ‘other’ variables was to a much lesser extent. Flowers were observed to warm ‘passively’ in pace with the environment, from the combined responses to environmental temperature and humidity. A result consistent with heat transfer between the flower and its surroundings. Furthermore, as expected elevated wind speed reduced floral temperatures, cooling flowers. In addition to these expected relationships, our study identifies that these ‘other’ weather variables can also influence temperature pattern contrasts, alongside light as the main determinant. For example, we found representative examples of certain locations of flowers varying in temperature from each other, beyond the effect of solar radiation (those indicated with patt* in figures 3-6). However, the effects of these ‘other variables’ and illumination on floral temperature do not occur in isolation, so these effects must be considered alongside the warming of flowers by illumination. Several of these effects appear to be representative of certain locations tending to be hotter, due to illumination effects. Therefore, they are affected more, relatively, by variables with negative influences. Wind speed had a greater effect on positions that tend to heat up more across species. Such positions, by virtue of being hotter allow greater heat transfer, and consequently a greater wind speed effect. Similarly, cloud cover affected *Coreopsis* reproductive structures more, paralleling its position-dependent illumination responses. Other position-dependent weather effects appear to indicate the increased importance of these variables to floral temperature at flower positions less influenced by illumination. In all species environmental temperature had a greater influence on positions, such as petal middle and edges, that are less influenced by illumination. Similarly, in *Cistus* ‘snow white’ we observed that relative humidity had a greater effect on such positions less affected by illumination.

The reduced importance of atmospheric pressure on the floral temperature and temperature patterns of the species monitored here may reflect that pressure would likely only influence floral temperature indirectly. That is, environment pressure may influence environmental wind speed, humidity and temperatures, thereby influencing floral temperature but these weather variables are already present in the models and have more direct effects. This may result in the remaining influence of pressure being small. Importantly, however, atmospheric pressure did not vary greatly within the study site (see table S9 Supplementary material), and thus interpretation is difficult. Perhaps variation in pressure would be greater between different sites of a flower species’ range, particularly with differing altitude, or time of year or across different environments.

### Remaining time of day effects

Following consideration of weather variables, Time of day was important in all species. The highest floral temperatures were observed in the early afternoon for each species. Contrast between flower positions was also influenced to a small extent by time of day in all species (figures 3-6b). This is most likely the effect of accumulated heat from prolonged exposure to solar radiation (the main source of warming of these flower species, see above). This demonstrates a need to consider not only the conditions at a given instant when investigating floral temperature generation but perhaps also the conditions prior to sampling, as heat can accumulate over time. Nevertheless, other factors may be involved in this relationship between time of day and floral temperature. Given that environmental temperature typically rises in the morning plateauing from midday until late evening (see Chow and Levermore, 2007; Peters and Evett, 2004; Reicosky et al., 1989), the observed time of day trend potentially reflects also daily environmental temperature trends. Plants also show daily cycles in transpiration activity, sometimes avoiding active stomatal transpiration where risk of water loss is highest during the hottest, driest parts of the day (Schulze and Hall, 1982; Trejo and Davis, 1991; Schroeder et al., 2001; Hetherington and Woodward, 2003; Lawson and Blatt, 2014; Simon et al., 2020). In this way flowers may be transpiring less later in the day, limiting heat loss and leading to higher floral temperature at certain parts of the day. It is not known if all four species sampled here possess active floral stomata or control transpiration activity in this way (see Cavallini-Speisser et al. (2021), but Harrap et al. (2020a) found *Coreopsis verticillata* to lack floral stomata). Further investigation should also consider influences of transpiration in flower daily cycles and possibly plant metabolic activity which may help explain the changes in floral temperature with time of day.

### Stability of floral temperature traits with the weather

Floral temperature monitoring took place during the day, across peak flowering of each flower species and under conditions where pollinators were active. Thus, the weather conditions and the changes in floral temperatures that occur alongside them represent those that would naturally occur while these flowers are interacting with pollinators. Our study finds the exact levels of illumination and other conditions, to lesser but varying degrees, realised by the flower at a given time will impact how elevated floral temperature is and how contrasting temperature patterns are. Additionally, how frequently these conditions co-occur will determine how frequently given floral temperatures are presented by flowers. This shifting in floral temperatures realised by the flower might have important consequences to the plant and the organisms they interact with and ultimately how much and how often floral temperature affects plant biology. Plants may experience variable susceptibility to microorganisms (Hildebrand et al., 2001; Rougerie-Durocher et al., 2020; Williamson et al., 1995, 2007), or viability of pollen, ovules and seeds (Mu et al., 2017; Hinojosa et al., 2019) with such changes in floral temperature. Metabolic activity of the flower may be similarly variable (Borghi et al., 2019, 2017; Borghi and Fernie, 2017).

The observed variations in floral temperature with weather conditions during pollinator activity may have particular effects on plant-pollinator interactions where such changes may alter the composition or salience of a temperature cue used by pollinators or pollinator preferences, therefore influencing pollinator responses to the flowers and potentially the fitness of both mutualists. Such dynamic floral temperature changes with the weather may be particularly relevant to the role of floral temperature as a pollinator learning or flower recognition cue, as conditions may change this signal to a state different from that the pollinator experienced and learnt previously. Such disruptions of other floral signalling modalities have been shown to impair subsequent recognition and learning of floral displays, however the presence of other signalling modalities can make floral displays more robust to such changes (Dyer and Chittka, 2004; Kaczorowski et al., 2012; Lawson et al., 2017). To explore these questions further, more studies need to address pollinator behavioural detection thresholds and the effect of shifting salience of floral display traits to understand what effect these shifts in temperature alongside the weather have.

While floral temperature is changeable with various weather conditions, floral temperature and temperature patterns were primarily dependent on illumination, and only moderate amounts of illumination (>300 lux/100) were required to achieve elevated floral temperatures relative to the environment and within flower temperature contrasts of the ∼2°C needed to elicit most pollinator responses. Furthermore, the shape of temperature patterns shown by flowers were not seen to shift with weather conditions. That is, while temperature contrast between flower positions changed with conditions, which positions of the flower where hotter or cooler than others did not (see figure 3-6). This meant, although the extent of elevated temperatures and salience of temperature cues will vary with conditions, elevated floral temperatures and contrasting temperature patterns should be present on flowers and are likely to be detectable by pollinators potentially influencing responses, across the vast majority of the conditions experienced during sampling. This suggests that, while variable, floral temperature traits persist to some degree over varied weather conditions as long as moderate amounts of illumination (and hence solar warming), are available. However, the dependence on at least some illumination to increase floral temperature may mean flowers in certain environments, those prone to less illumination, may not be able to utilise floral temperature to maintain pollen and ovule viability or signal to pollinators, unless heat is produced by alternative means such as *via* thermogenesis. Further investigation of the use of temperature cues by pollinators foraging in such environments as well as the incidence of traits that may encourage floral warming with illumination (discussed above) or other adaptations that may mitigate the inability to warm in low illumination and across different habitats may expand our understanding of floral temperatures’ utilisation.

## Acknowledgements

The authors would like to thank the National Botanic Garden of Wales and their staff for use of their facilities and their cooperation with this research project. The authors would also like to thank the Met Office (UK) National Meteorological library and archive for providing regional weather data for this research project.

MJMH was supported by a Natural Environment Research Council studentship within the GW4 + Doctoral Training Partnership (studentship NE/L002434/) and a Bristol Centre for Agricultural Innovation grant awarded to SAR. HMW was supported by the Biotechnology & Biological Sciences Research Council (grant BB/M002780/1). The funding bodies played no role in the study design, data collection, analysis, interpretation or the writing of this manuscript.

## Author contributions

MJMH was involved in investigation, data collection, formal analysis, data visualization, writing and implementation of code, data curation and wrote the original draft of this manuscript. MJMH is also the ‘thermographer’ indicated in the text, possessing an Infrared Training Centre Level 1 Thermographer qualification. Resources and facilities for this research were obtained by NdV, NHdI, HMW, and SAR. Funding acquisition as well as project supervision and administration were conducted by NdV, NHdI, HMW, SAR. All Authors took part in project conceptualization, methodological design and writing, review and editing. All authors approved the final version of the manuscript.

## Data availability statement

The thermal images collected within this research are freely available on the Zenodo data repository (Harrap et al., 2024b). Floral temperature data, weather data and code associated with the analyses of floral temperature changes with weather conditions, correlation of weather conditions and comparison of weather conditions between flower species, as well as code for figure generation is freely available on the Zenodo data repository (Harrap et al., 2024a).

Weather data provided by the Pembrey Sands weather station from 2016 and 2017 was obtained from the Met Office (UK) National Meteorological library and archive for this research project. The following copyright statement applies to this weather data:

© Crown Copyright 2016 and 2017. Information provided by the National Meteorological Library and Archive – Met Office, UK.

