## Supplementary Materials for "Variations of floral temperature in changing weather conditions"

**Supplementary Information 1:** *Details of thermographs used in Figure 2*

Below we list further details about the specific thermographs used in figure 2 and the weather conditions at time of thermograph capture. 'Individual ID number' noted below correspond with the 'Global ID' in datafiles associated with this publication (Harrap et al., 2024), 'observation' refers to the repeat thermograph measurements conducted on each individual flower (first observation being 1, second being 2, *etc*.). All other variables, and how they are assigned and collected, are as described elsewhere in the text.

*Cistus* 'snow fire': individual ID number *175f2017y* observation 3; Time of thermograph (DD.M.YY at HH.MM) = 15.8.17 at 12.29; Illumination = 1201 Lux/100; hourly environmental temperature = 17.4 ºC; hourly mean windspeed = 16 kn; hourly total cloud cover = 0 oktas; hourly relative humidity = 77.9%; hourly pressure at mean sea level= 1016 hPa.

*Cistus* 'snow white', individual ID number *143f2017y* observation 3, Time of thermograph (DD.M.YY at HH.MM) = 15.8.17 at 12.33; Illumination = 1163 Lux/100; all other conditions as *Cistus* 'snow fire'

*Coreopsis verticillata*, individual ID number *45f2017y* observation 10, Time of thermograph (DD.M.YY at HH.MM)= 25.7.17 at 12.46; Illumination = 1169 Lux/100; hourly environmental temperature = 19.5 ºC; hourly mean windspeed = 6 kn; hourly total cloud cover = 0 oktas; hourly relative humidity = 82%; hourly pressure at mean sea level= 1017 hPa.

*Geranium psilostemon*, individual ID number *40f2017y* observation 13, Time of thermograph (DD.M.Y at HH.MM)= 14.7.17 at 15.42; Illumination = 1098 Lux/100; hourly environmental temperature = 19 ºC; hourly mean windspeed = 13 kn; hourly total cloud cover = 6 oktas; hourly relative humidity = 61.9%; hourly pressure at mean sea level= 1023 hPa.

**Supplementary Information 2:** *Assessment of weather variable correlation*

The overall weather conditions experienced by flowers at any one time are a combination of the many weather variables measured. A flower will not experience one variable (*e.g.* illumination) in complete absence of another (*e.g.* environmental temperature). Weather conditions themselves are likely to correlate, as they often influence each other. Consequentially, before assessment of weather influences on floral temperatures, it is important to understand the relationships between various weather variables. Furthermore, although the methods used for our analysis of weather influences on floral temperatures (AIC based modelling) is robust to moderate covariation between variables ($-0.5 < r < 0.5$, Freckleton, 2011) it is important to explore the covariation between weather variables and identify covariation that may conflate our analyses.

A complication to exploring correlation between weather variables is that different weather variables used are measured at different timescales. That is, illumination (Lux/100) is measured multiple times per hour at every thermography incident, while all other variables are measured hourly (see methods). It would be inappropriate to assess correlation between variables measured hourly at a thermography incident scale, as correlations would likely reflect thermography sampling effort, the hours where more thermography incidents took place. Consequentially, the correlation between the variables collected hourly at each hour during which thermographs were collected (*i.e.* every unique hour where we take thermal images goes into this dataset once - to a total of 340 hours across both years) were compared against each other (15 comparisons). Then correlation between each unique thermography incident's illumination (Lux/100) measurements (2999 instances) and the hourly data variables was compared (6 comparisons). All correlations were assessed in *R* 3.6.3 (R Core Team, 2020) with Spearman's rank correlation tests using the *rcorr* function in the *R* package *Hmisc* 4.4-1 (Harrell Jr et al., 2020). This test was favoured as many weather variables (particularly environmental temperature, hour of the day, cloud cover) did not appear to show normal distributions. Following this, a further analysis was conducted assessing how each weather variable differed with hourly mean wind direction. This analysis, as before, was conducted at the hours at which thermography took place or thermography incident scales for variables measured hourly and illumination respectively (7 more comparisons). Hourly mean wind direction was provided in degrees by the Pemberley Sands weather station, this was converted to a categorical variable for eight compass point directions (N., N.E., E., S.E., S., S.W., W. and N.W.). For full classification of hourly mean wind direction by compass point directions, see the code associated with this publication (Harrap et al., 2024). The relationship between this hourly mean wind direction and each weather variable was assessed in *R* 3.6.3 using Kruskal-Wallis tests. *Post hoc* comparisons between groups were carried out with Dunn tests, using the *R* package *FSA* version 0.8.30 (Ogle et al., 2020). To account for the effects of multiple comparisons conducted within this assessment of correlation between our weather variables a Bonferroni correction was applied to this analysis, with the new P criterion being 0.0018 (0.05 / [15+6+7]).

Several correlations between hourly weather variables and illumination and other weather variables were identified (figure S1). However the strength of these correlations was moderate ($-0.5 < r < 0.5$) to weak ($-0.3 < r < 0.3$) and therefore unlikely to interfere with AIC model comparison's ability to resolve the influences of each weather variables on floral temperature in our later analysis (Freckleton, 2011). Consequently, exclusion of these variables or creation of composite parameters was deemed unnecessary. Correlations observed between variables measured hourly (figure S1A) make logical sense when we consider warming of the environment over the course of the day (hour of the day with environmental temperature), the effects of air motion (wind speed with environmental temperature), evaporation rates (relative humidity and temperature) and the water cycle (relative humidity with cloud cover) on different weather conditions. Illumination (Lux/100) correlated with all hourly weather variables in some way (figure S1B). However, several of these correlations were weak. The strongest of these relationships between illumination and hourly variables continued to make logical sense when we consider how solar radiation warms the environment (illumination with environmental temperature and humidity) and particularly the effect of cloud cover on light conditions (illumination with cloud cover). That hourly weather variables and illumination correlated logically, even though hourly variables were measured at Pemberley Sands 21.4km away from the Carmarthen study site, indicates that these variables measured hourly are still at least indicatory of the corresponding weather conditions at the Carmarthen study site.

Hourly mean wind direction (expressed as the prevailing compass point direction see above) had a significant effect on all weather variables (table S1 for Kruskal-Wallis test results and figure S2). All weather conditions varied with wind direction. *Post hoc* tests suggested there were two groupings of wind directions that coincided with similar conditions: the first group comprising of North, North-eastern and South-eastern wind directions; the second of all other wind directions. That hourly mean wind direction corresponds with other variables means that changes to conditions that occur with changes in wind direction will likely be described by other weather variables. Furthermore, exactly what conditions correspond with specific wind directions will vary greatly from location to location. Consequently, it was deemed more informative to not include wind direction in our further analysis, whose effects are likely region-specific or reflect changes in other conditions with wind direction. Additionally, this exclusion reduced the complexity of statistical analyses.

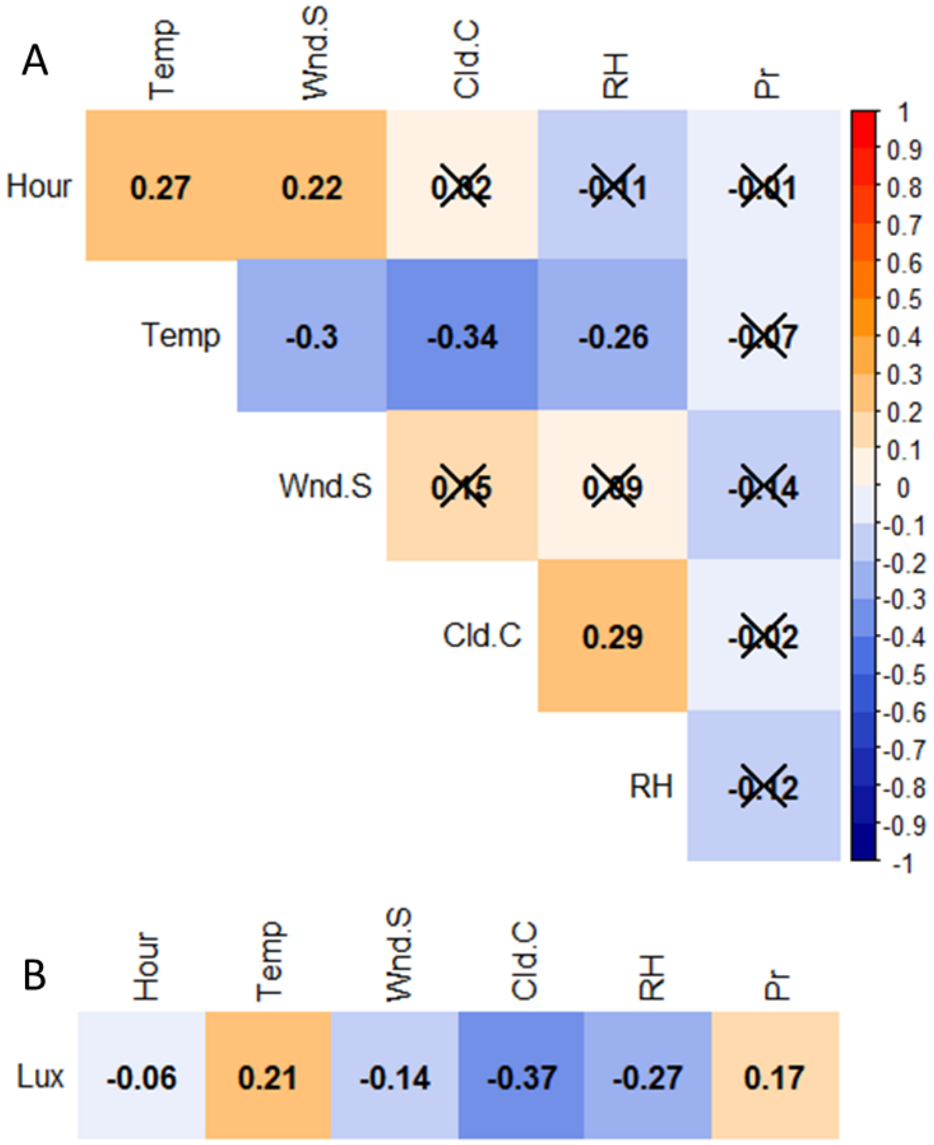

**Figure S1:** Correlation matrixes comparing the correlation between **A** each weather variable measured hourly, and **B** Illumination and weather variables measured hourly. Weather variable labelling is as follows: 'Hour', hour of the day; 'Temp', environmental temperature (°c); 'Wnd.S', wind speed (kn); 'Cld.C', cloud cover (oktas); 'RH', relative humidity (%); 'Pr', atmospheric pressure (hPa); and 'Lux', illumination (lux/100, see main text). Strength and direction of correlation between variable (*r*) is indicated by the value within the matrices and the colour of each box according to the scale to the right of panel **A**. Crossed-out comparisons indicate non-significant correlations (*p* criterion for significance = 0.0018).

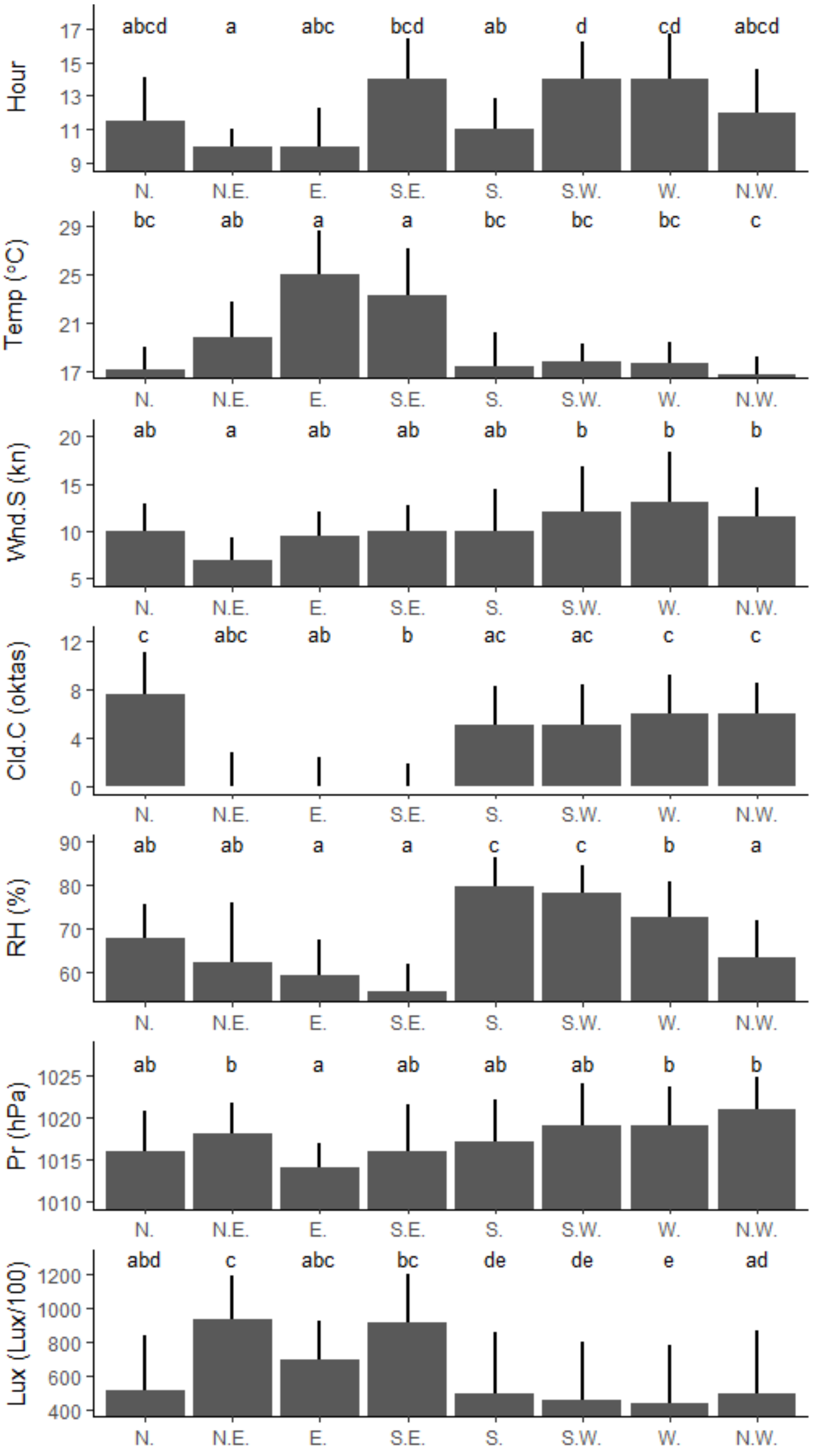

**Figure S2:** How each weather variable changes with hourly mean wind direction (expressed as the prevailing compass point direction). Panel x axis labels indicate the weather variables as follows: 'Hour', hour of the day; 'Temp', environmental temperature (°c); 'Wnd.S', wind speed (kn); 'Cld.C', cloud cover (oktas); 'RH', relative humidity (%); 'Pr', atmospheric pressure (hPa); and 'Lux', illumination (lux/100, see main text). Plotted are the median values of each weather variable during hours (variables measured hourly) and thermography incidents (Lux) where hourly mean wind direction was each compass direction. Error Bars represent plus one standard deviation. Letters above bars indicate groupings according to Dunn tests, wind directions sharing a letter indicating that different wind directions are not significantly different for each weather variable ($p$ criterion for significance = 0.0018).

**Table S1:** Results of Kruskal-Wallis analyses of how hourly mean wind direction corresponds with other weather variables. Given are the summary statistics (Chi-squared, df, *p*) of Kruskal-Wallis tests comparing median values of each weather variable between hours (hourly variables) and thermography incidents (illumination) where mean hourly wind direction was in each compass point direction (*p* criterion for significance = 0.0018).

| Weather Variable | Chi-squared | df | *p* |
|---:|---:|---:|---:|
| Hour of the day | 37.6 | 7 | <0.0001 |
| Environmental temperature | 76.3 | 7 | <0.0001 |
| Wind Speed | 26.5 | 7 | 0.0004 |
| Cloud cover | 52.4 | 7 | <0.0001 |
| Relative Humidity | 157.9 | 7 | <0.0001 |
| Atmospheric Pressure | 27.6 | 7 | 0.0003 |
| Illumination | 136.6 | 7 | <0.0001 |

**Supplementary Information 3:** *Comparison of weather variables experienced by flower species*

The different flower species were not sampled simultaneously (table 2) and may differ in the conditions in which they were thermographed. This is the case even with the two *Cistus* varieties, that were sampled during the same period due to the thermographer being unable to image more than one flower at the exact same instant. This may lead to erroneous conclusions that a weather variable has differing influences on floral temperature of different species, when this is simply due to a difference in the range of that weather variable experienced by each species. If we are to compare across the flower species which weather variables are important and compare the nature (direction, effects size, influences on temperature patterns *etc.*) of weather influences to identify common trends in floral temperature responses, it is important to understand the extent that the different flower species differ in the conditions experienced. Consequentially the values of each of the weather variables used in our analysis of floral temperature (not including wind direction) at each thermography incident were compared between species using Kruskal-Wallis tests, with *post hoc* Dunn tests. Again, a Bonferroni correction was applied to this analysis, with the $p$ criterion being calculated as 0.007 (= 0.05/7). For these analyses hour of the day was expressed as hour of the day after 9am (hour of the day - 9) and atmospheric pressure was expressed as difference in hPa from 'normal' conditions (ΔhPa). Where normal atmospheric pressure conditions were defined as 1013 hPa. These transformations of the parameters are applied within our models and discussed further in Supplementary material 4.

Species differed significantly in conditions experienced across all weather variables (table S2). *Post hoc* testing revealed that the two *Cistus* varieties tended to experience similar conditions. This was unsurprising as these two varieties were sampled during the same periods of time. Although the two *Cistus* varieties differed in cloud cover conditions experienced. *C. verticillata* experienced similar conditions to the *Cistus* varieties in terms of illumination and pressure but differed in other respects. *G. psilostomon* experienced conditions that were more similar to *C. verticillata* then to the *Cistus* varieties but differed in several variables. Although significant variation in conditions experienced by species existed, the average conditions and variation about these, as well as the range of conditions experienced by each species (see table S2) were still largely similar. Consequently, it was deemed that species experienced comparable, although differing, weather conditions and it was unlikely that differences in the conditions experienced by species will conflate the identification of the relative influences of each weather condition on floral temperature.

**Table S2:** Results of Kruskal-Wallis analyses comparing conditions experienced by each flower species across all thermography incidents. Given for each variable are the summary statistics (Chi-squared, df, p) of Kruskal-Wallis tests comparing median values of each weather variable between species. Additionally, within sub tables are summary values (Median, SD, n, min and max) of each weather variable experienced by each species. Column letter indicates grouping according to *post hoc* Dunn tests. Species sharing a letter indicates that species are not significantly different in conditions experienced for that weather variable (p criterion for significance = 0.007).

**Hour of the day (after 9am),** (Kruskal-Wallis tests: Chi-squared=66.7 df=3 $p$<0.001)

| Species | Median | SD | n | min | max | Letter |
|---|---|---|---|---|---|---|
| *C. verticillata* | 5 | 2.6 | 837 | 0 | 9 | a |
| *Cistus* 'snow fire' | 3 | 2.3 | 1060 | 0 | 8 | b |
| *Cistus* 'snow white' | 3 | 2.2 | 926 | 0 | 8 | b |
| *G. psilostomon* | 3 | 2.4 | 901 | 0 | 9 | a |

**Environmental temperature (°C)** (Kruskal-Wallis tests: Chi-squared=371.4 df=3 $p$<0.001)

| Species | Median | SD | n | min | max | Letter |
|---|---|---|---|---|---|---|
| *C. verticillata* | 18.7 | 2.2 | 837 | 14.1 | 24.0 | a |
| *Cistus* 'snow fire' | 17.2 | 2.8 | 1060 | 13.9 | 27.9 | b |
| *Cistus* 'snow white' | 17 | 2.8 | 926 | 13.9 | 27.9 | b |
| *G. psilostomon* | 18.3 | 2.9 | 901 | 14.5 | 30.4 | a |

**Wind Speed (kn)** (Kruskal-Wallis tests: Chi-squared=60.3 df=3 $p$<0.001)

| Species | Median | SD | n | min | max | Letter |
|---|---|---|---|---|---|---|
| *C. verticillata* | 11 | 4.4 | 837 | 3 | 24 | a |
| *Cistus* 'snow fire' | 11 | 4.9 | 1060 | 3 | 22 | b |
| *Cistus* 'snow white' | 11 | 4.9 | 926 | 3 | 22 | b |
| *G. psilostomon* | 11 | 3.8 | 901 | 2 | 20 | a |

**Cloud cover (oktas)** (Kruskal-Wallis tests: Chi-squared=224.3 df=3 $p$<0.001)

| Species | Median | SD | n | min | max | Letter |
|---|---|---|---|---|---|---|
| *C. verticillata* | 5 | 3.1 | 837 | 0 | 8 | a |
| *Cistus* 'snow fire' | 3 | 3.2 | 1060 | 0 | 8 | b |
| *Cistus* 'snow white' | 1 | 3.2 | 926 | 0 | 8 | c |
| *G. psilostomon* | 7 | 3.3 | 901 | 0 | 8 | d |

**Relative Humidity (%)** (Kruskal-Wallis tests: Chi-squared=84.1 df=3 $p$<0.001)

| Species | Median | SD | n | min | max | Letter |
|---|---|---|---|---|---|---|
| *C. verticillata* | 68.6 | 9.8 | 837 | 48.9 | 90.6 | a |
| *Cistus* 'snow fire' | 74.4 | 8.0 | 1060 | 50.8 | 93.3 | b |
| *Cistus* 'snow white' | 75.2 | 7.5 | 926 | 50.8 | 89.8 | b |
| *G. psilostomon* | 71.3 | 12.5 | 901 | 47.3 | 100.0 | c |

**Atmospheric Pressure (ΔhPa)** (Kruskal-Wallis tests: Chi-squared=32.8 df=3 $p$<0.001)

| Species | Median | SD | n | min | max | Letter |
|---|---|---|---|---|---|---|
| *C. verticillata* | 5 | 6.4 | 837 | -8 | 14 | a |
| *Cistus* 'snow fire' | 6 | 3.7 | 1060 | -5 | 11 | a |
| *Cistus* 'snow white' | 6 | 3.6 | 926 | -5 | 11 | a |
| *G. psilostomon* | 7 | 4.6 | 901 | -5 | 14 | b |

**Illumination (lux/100),** Kruskal-Wallis tests: Chi-squared=32.3 df=3 $p$<0.001)

| Species | Median | SD | n | min | max | Letter |
|---|---|---|---|---|---|---|
| *C. verticillata* | 436.0 | 337.7 | 837 | 78 | 1343 | a |
| *Cistus* 'snow fire' | 462.5 | 371.8 | 1060 | 32 | 1385 | a |
| *Cistus* 'snow white' | 462.0 | 371.7 | 926 | 27 | 1390 | a |
| *G. psilostomon* | 550.0 | 321.1 | 901 | 112 | 1401 | b |

**Supplementary information 4:** *Floral temperature models and simplification procedures*

The ‘full model’ of weather variable and time of day effects on floral temperature and temperature patterns fitted to each species of flowers, before any model simplification was applied, is as follows:

$$\begin{aligned} Y_{fne} = {} & i_a + B(i_b) + C(i_c) + D(i_d) \\ & + (H-9) \times \big(h_a + B(h_b) + C(h_c) + D(h_d)\big) \\ & + (H-9)^2 \times \big(j_a + B(j_b) + C(j_c) + D(j_d)\big) \\ & + \ln(L) \times \big(l_a + B(l_b) + C(l_c) + D(l_d)\big) \\ & + T \times \big(t_a + B(t_b) + C(t_c) + D(t_d)\big) \\ & + O \times \big(o_a + B(o_b) + C(o_c) + D(o_d)\big) \\ & + R \times \big(r_a + B(r_b) + C(r_c) + D(r_d)\big) \\ & + (P-1013) \times \big(p_a + B(p_b) + C(p_c) + D(p_d)\big) \\ & + W \times \big(w_a + B(w_b) + C(w_c) + D(w_d)\big) + q_n \,, \end{aligned} \qquad \text{(S1)}$$

where $Y_{fne}$ is the temperature in °C of position $f$ on flower $n$ during thermography incident $e$. The flower positions ($f$) being those measured during our sampling (a, reproductive structures; b, petal base; c, petal middle; d, petal tip). Parameter $i_a$, the model intercept, is the floral temperature of the reproductive structures of the given species independent of effects of weather variables or time of day. this model intercept can be altered depending on position of a temperature measurement on the flower (flower position). Parameters $i_b$, $i_c$, and $i_d$ describe the difference in temperature independent of weather variable and time of day effects (the model intercept) between the reproductive structures of the flower and the petal base, petal middle and petal tip respectively. The application of parameters $i_b$, $i_c$, and $i_d$ are controlled by Boolean operators $B$, $C$ and $D$ respectively depending on the flower position being measured ($f$), where:

$$B = \begin{cases} 0, & flower\ position\ is\ not\ the\ petal\ base \\ 1, & flower\ position\ is\ the\ petal\ base \end{cases}, \qquad \text{(S2)}$$

$$C = \begin{cases} 0, & flower\ position\ is\ not\ the\ petal\ middle \\ 1, & flower\ position\ is\ the\ petal\ middle \end{cases}, \qquad \text{(S3)}$$

and

$$D = \begin{cases} 0, & flower\ position\ is\ not\ the\ petal\ edge \\ 1, & flower\ position\ is\ the\ petal\ edge \end{cases}. \qquad \text{(S4)}$$

The actions of Boolean operators $B$, $C$ and $D$ alter the model intercept depending on the flower position being measured. When Boolean operators $B$, $C$ and $D$ have nonzero values parameters $i_b$, $i_c$, and $i_d$ are added to $i_a$. When Boolean operators $B$, $C$ and $D$ have values of zero, such as when the reproductive structures flower position is being measured no value is added to $i_a$ and the model intercept is described by $i_a$ alone. It is important to note (as it is the basis for the tests of effects on temperature patterns during model simplification) that when parameters $i_b$, $i_c$, and $i_d$ have values of zero the model intercept is likewise described by $i_a$ alone. These same Boolean operators control other flower position-dependent effects, such as flower position-dependent effects of weather variables, in the same manner as they control the position-dependent model intercepts. Flower identity was included as a random factor, $q_n$, which represents the change in floral temperature, and thus model intercept, relative to $i_a$ for individual flower *n*.

The hour of the day in which thermography incident $e$ took place is given by parameter $H$. In the model this is transformed to "hour of the day after 9:00" in which thermography incident $e$ took place by subtracting 9 from this value. This was done to aid fitting of parameter values by the statistical software and so that the values of the weather variables start at a point approximately zero, like other variables, making comparisons between parameter values of models that have and lack these parameter's effects more straightforward. Parameters $h_a$ and $j_a$ describe the change in reproductive structure floral temperature with time of day, together allowing a quadratic relationship. Parameters $h_a$ and $j_a$ are modified at other flower positions. Parameters $h_b$, $h_c$, and $h_d$ describe the time-of-day effects relative to $h_a$, between the reproductive structures of the flower and the petal base, petal middle and petal tip respectively. Parameters $j_b$, $j_c$, and $j_d$ describe the time-of-day effects relative to $j_a$, between the reproductive structures of the flower and the petal base, petal middle and petal tip respectively. Boolean operators $B$, $C$ and $D$ control application of both $h_b$, $h_c$, $h_d$, $j_b$, $j_c$, and $j_d$ dependent on the flower position measured as described above (equations S2-4).

Illumination (in Lux/100) at the time of thermography incident $e$ is given by $L$, this is log-transformed in the model. The change in floral temperature of reproductive structures with increased illumination is described by $l_a$. The model allows other flower positions to differ in their responses to illumination. The difference in changes in floral temperature with increased illumination of other flower positions relative to $l_a$ are described by $l_b$, $l_c$, and $l_d$ for the petal base, petal middle and petal tip respectively. Boolean operators $B$, $C$ and $D$ control application of both $l_b$, $l_c$, and $l_d$ dependent on the flower position measured in the same way as described above (equations S2-4).

Environmental temperature (in °C) at the time of thermography incident $e$ is given by $T$. The change in floral temperature of reproductive structures with increased environmental temperature is described by $t_a$. The model allows other flower positions to differ in their responses to environmental temperature. The difference in changes in floral temperature with increased environmental temperature of other flower positions relative to $t_a$ are described by $t_b$, $t_c$, and $t_d$ for the petal base, petal middle and petal tip respectively. Boolean operators $B$, $C$ and $D$ control application of $t_b$, $t_c$, and $t_d$ dependent on the flower position measured in the same way as described above (equations S2-4).

Cloud cover (in oktas) at the time of thermography incident $e$ is given by $O$. The change in floral temperature of reproductive structures with increased cloud cover is described by $o_a$. The model allows other flower positions to differ in their responses to cloud cover. The difference in changes in floral temperature with increased cloud cover of other flower positions relative to $o_a$ are described by $o_b$, $o_c$, and $o_d$ for the petal base, petal middle and petal tip respectively. Boolean operators $B$, $C$ and $D$ control application of $o_b$, $o_c$, and $o_d$ dependent on the flower position measured in the same way as described above (equations S2-4).

Percentage relative humidity at the time of thermography incident $e$ is given by $R$. The change in floral temperature of reproductive structures with increased humidity is described by $r_a$. The model allows other flower positions to differ in their responses to relative humidity. The difference in changes in floral temperature with increased environmental temperature of other flower positions relative to $r_a$ are described by $r_b$, $r_c$, and $r_d$ for the petal base, petal middle and petal tip respectively. Boolean operators $B$, $C$ and $D$ control application of $r_b$, $r_c$, and $r_d$ dependent on the flower position measured in the same way as described above (equations S2-4).

Atmospheric pressure (in hPa) at the time of thermography incident $e$ is given by $P$. This is transformed in the model to be difference from 'normal' atmospheric pressure (ΔhPa) by subtracting 1013 ('normal' atmospheric pressure) from $P$. This transformation was done for the exact same reasons as the transformation applied to $H$. The change in floral temperature of reproductive structures with atmospheric pressure is described by $p_a$. The model allows other flower positions to differ in their responses to pressure. The difference in changes in floral temperature with increased atmospheric pressure of other flower positions relative to $p_a$ are described by $p_b$, $p_c$, and $p_d$ for the petal base, petal middle and petal tip respectively. Boolean operators $B$, $C$ and $D$ control application of $p_b$, $p_c$, and $p_d$ dependent on the flower position measured in the same way as described above (equations S2-4).

Average wind speed (in kn) at the time of thermography incident $e$ is given by $W$. The change in floral temperature of reproductive structures with increased wind speed is described by $w_a$. The model allows other flower positions to differ in their responses to wind speed. The difference in changes in floral temperature with increased wind speed of other flower positions relative to $w_a$ are described by $w_b$, $w_c$, and $w_d$ for the petal base, petal middle and petal tip respectively. Boolean operators $B$, $C$ and $D$ control application of $w_b$, $w_c$, and $w_d$ dependent on the flower position measured in the same way as described above (equations S2-4).

As described in the main text, the effects on floral temperature of each weather variable, hour of day and how these effects and floral temperature vary with positions on the flower was assessed for each species independently using AIC model simplification techniques in *R* version 3.6.3 (R Core Team, 2020), utilizing the package *lme4* 1.1.-25 (Bates et al., 2015) and AIC comparison criteria laid out by Richards (2008). This involved paired AIC comparisons between a standing best model and a simpler model constructed by removing parameters from the standing best model (by forcing those parameters to be zero). If removal of parameters resulted in a sufficient increase in AIC the standing best (more complex) model would remain the best for the next comparison. If otherwise, the simpler model would become the standing best model for the next comparison. Initially the full model described in equation S1 was the initial standing best model for the first comparison. These comparisons followed a set sequence for each species testing the effects of each variable in turn (see table S3). For each variable the effects on contrast and structure of temperature patterns was assessed first (part a of each comparison in table S3). This was done by removing position-dependent effects relevant to the variable in question to produce a simpler model for comparison with the standing best model. Following this the absolute effects of that variable were tested by removing position-dependent effects (if retained after the previous comparison) and position-independent effects for a subsequent comparison (part b of each comparison in table S3). This sequence of comparisons is discussed further in the main text. The sequence of comparisons applied to each species and the parameters removed from the standing best model to make simpler models at each comparison is described in table S3.

**Table S3:** The sequence of paired comparisons used to assess the effects of weather variables and time of day on floral temperature and temperature patterns. The sequence of comparisons carried out is shown, in the order described by column '#'. These comprise of two parts where for each variable we test the position-dependent effects on floral temperature and then absolute effects on floral temperature. This involves fitting a simpler model generated by removing the parameters described at each comparison and comparing it to the standing best model of the previous comparison (starting with the full model described in equation S1). Note that during the test of the absolute effects, the parameters retained by the previous position-dependent effects test are removed again (see main text and supplementary Information 4). Additionally, note that absolute intercept effects and random factor effects, although included in the model, are not tested.

| # | Variable | Part | Effect tested | Parameters removed (if retained in model) |
|---|---|---|---|---|
| 1 | Wind speed (kn) | a | Position-dependent effect | $w_b, w_c, w_d$ |
| | | b | Absolute effect | $w_a, w_b, w_c, w_d$ |
| 2 | Pressure (ΔhPa) | a | Position-dependent effect | $p_b, p_c, p_d$ |
| | | b | Absolute effect | $p_a, p_b, p_c, p_d$ |
| 3 | Relative Humidity (%) | a | Position-dependent effect | $r_b, r_c, r_d$ |
| | | b | Absolute effect | $r_a, r_b, r_c, r_d$ |
| 4 | Cloud Cover (oktas) | a | Position-dependent effect | $o_b, o_c, o_d$ |
| | | b | Absolute effect | $o_a, o_b, o_c, o_d$ |
| 5 | Environmental temperature (°C) | a | Position-dependent effect | $t_b, t_c, t_d$ |
| | | b | Absolute effect | $t_a, t_b, t_c, t_d$ |
| 6 | Illumination (ln[Lux/100]) | a | Position-dependent effect | $l_b, l_c, l_d$ |
| | | b | Absolute effect | $l_a, l_b, l_c, l_d$ |
| 7 | Hour of day (after 9am) | a | Position-dependent effect | $h_b, h_c, h_d, j_b, j_c, j_d$ |
| | | b | Absolute effect | $h_a, h_b, h_c, h_d, j_a, j_b, j_c, j_d$ |
| 8 | Intercept | a | Position-dependent effect | $i_b, i_c, i_d$ |

Following the paired comparisons described in table S3, a further set of comparisons was carried out to assess whether the different flower positions differ in their responses to weather conditions and floral temperature. Alternative versions of the standing best model (maintaining the same model structure and effects) were constructed where different flower positions were grouped together. These alternative flower position models effectively treated measurements of floral temperature at flower positions grouped together as a repeat measurement of a single flower position. This was achieved by altering the inclusion and operation of Booleans $B$, $C$ and $D$. The removal of the Boolean operator relevant to each flower position ($B$, for the petal base; $C$ for the petal middle and $D$ for the petal edge) with no other adjustments to other Booleans would make that flower position equivalent to the flower's reproductive structures (see equation S1). Removal of the relevant Boolean and adjustments of others to include measurements taken from that position allows other positions to be made equivalent in these alternative models. For example, to make a model where the petal base and petal middle equivalent but the reproductive structures and petal edge remain distinct from these positions and each other (model Ap12 – see code associated with this publication and table S8 below) Boolean $C$ would be removed (set to zero) and $B$ would be altered as follows:

$$B = \begin{cases} 0, flower\ position\ is\ not\ the\ petal\ base\ or\ petal\ middle \\ 1, \quad flower\ position\ is\ the\ petal\ base\ or\ petal\ middle \end{cases}. \tag{S5}$$

These techniques were also applied to group more than one position together by removing several Booleans and/or altering Booleans further. For example, to make the petal base, petal middle and petal edge equivalent but distinct from the reproductive structures (such as in model Ap10 – see code associated with this publication and table S8 below) Booleans $C$ and $D$ would be removed (fixing their values to zero) and $B$ would be altered as follows:

$$B = \begin{cases} 0, flower\ position\ is\ reproductive\ strucutres \\ 1, flower\ position\ not\ reproductive\ structures \end{cases}. \tag{S6}$$

Using such techniques 15 alternative versions of the standing best model produced by the paired comparisons with different grouping structures of different flower positions were generated and fit to the data. These 15 models included a model where all positions were treated as equivalent to each other, the 'Absent model', where position-dependent effects were absent produced by removing Booleans $B$, $C$ and $D$. These 15 models also included the standing best model at the end of the paired comparisons described in table S3, the 'standing best' where all four positions differed from each other. The remaining 13 alternative position models 'Ap models' included all the remaining grouping structures of the four flower positions. The position grouping structures of each model are described in table S8 and in the code associated with this publication. These alternative position models were compared with AIC. The best fitting model at the end of this AIC comparison was considered the best model of floral temperature for each species.

**Supplementary Table S4:** The results of model selection testing weather effects on *Cistus* 'snow fire'. Comparisons of standing best models and a simpler version where parameters are removed are given for each effect tested in our model selection process. Bold AIC value indicates best performing model of each comparison. AIC is given for both models (note how standing best model AIC matches one of the previous model AICs). Also given is an assessment of model fit, Δdeviance=Δdev. A verdict on each comparison is given: Asterisks '∗' indicate parameters where AIC comparisons indicate inclusion of effects in best model, based on Richards (2008).

| Tested effect | | Standing best model AIC | Simpler model AIC | ΔAIC | Δdev. | df | *p* | Verdict |
|---|---|---|---|---|---|---|---|---|
| Wind speed (kn) | Position-dependent effect | **85841.99** | 86081.46 | 239.47 | 245.47 | 3 | <0.001 | ∗ |
| | Absolute effect | **85841.99** | 86195.03 | 353.04 (113.57) | 361.04 | 4 | <0.001 | ∗ |
| Pressure (ΔhPa) | Position-dependent effect | 85841.99 | **85839.52** | 2.47 | 3.53 | 3 | 0.317 | |
| | Absolute effect | **85839.52** | 85885.96 | 46.44 | 48.44 | 1 | <0.001 | ∗ |
| Relative Humidity (%) | Position-dependent effect | **85839.52** | 85846.55 | 7.03 | 13.04 | 3 | 0.005 | ∗ |
| | Absolute effect | **85839.52** | 85919.52 | 80.00 (72.97) | 88.00 | 4 | <0.001 | ∗ |
| Cloud Cover (oktas) | Position-dependent effect | 85839.52 | **85838.50** | 1.02 | 4.99 | 3 | 0.173 | |
| | Absolute effect | **85838.50** | 85869.26 | 30.76 | 32.76 | 1 | <0.001 | ∗ |
| Environmental temperature (°C) | Position-dependent effect | **85838.50** | 86226.42 | 387.92 | 393.92 | 3 | <0.001 | ∗ |
| | Absolute effect | **85838.50** | 86422.30 | 583.80 (195.88) | 591.79 | 4 | <0.001 | ∗ |
| Illumination (ln[Lux/100]) | Position-dependent effect | **85838.50** | 92340.20 | 6501.70 | 6507.70 | 3 | <0.001 | ∗ |
| | Absolute effect | **85838.50** | 98993.57 | 13155.07 (6653.37) | 13163 | 4 | <0.001 | ∗ |
| Hour of day (after 9am) | Position-dependent effect | **85838.50** | 86106.93 | 268.43 | 280.43 | 6 | <0.001 | ∗ |
| | Absolute effect | **85838.50** | 89654.76 | 3816.26 (3547.83) | 3832.3 | 8 | <0.001 | ∗ |
| Intercept | Position-dependent effect | **85838.50** | 86212.04 | 373.54 | 379.54 | 3 | <0.001 | ∗ |
| | Model intercept | | | | | | | - |

**Supplementary Table S5:** The results of model selection testing weather effects on *Cistus* 'snow white'. Comparisons of standing best models and a simpler version where parameters are removed are given for each effect tested in our model selection process. Bold AIC value indicates best performing model of each comparison. AIC is given for both models (note how standing best model AIC matches one of the previous model AICs). Also given is an assessment of model fit, Δdeviance=Δdev. A verdict on each comparison is given: Asterisks '∗' indicate parameters where AIC comparisons indicate inclusion of effects in best model, based on Richards (2008).

| Tested effect | | Standing best model AIC | Simpler model AIC | ΔAIC | Δdev. | df | $p$ | Verdict |
|---|---|---|---|---|---|---|---|---|
| Wind speed (kn) | Position-dependent effect | **63429.55** | 63565.58 | 136.03 | 142.02 | 3 | <0.001 | ∗ |
| | Absolute effect | **63429.55** | 63969.56 | 540.01 (403.98) | 548.01 | 4 | <0.001 | ∗ |
| Pressure (ΔhPa) | Position-dependent effect | 63429.55 | **63424.98** | 4.57 | 1.43 | 3 | 0.699 | |
| | Absolute effect | **63424.98** | 63470.15 | 45.17 | 47.16 | 1 | <0.001 | ∗ |
| Relative Humidity (%) | Position-dependent effect | **63424.98** | 63470.45 | 45.47 | 51.46 | 3 | <0.001 | ∗ |
| | Absolute effect | **63424.98** | 63517.16 | 92.18 (46.17) | 100.18 | 4 | <0.001 | ∗ |
| Cloud Cover (oktas) | Position-dependent effect | 63424.98 | **63419.33** | 5.65 | 0.35 | 3 | 0.951 | |
| | Absolute effect | **63419.33** | 63509.57 | 90.24 | 92.24 | 1 | <0.001 | ∗ |
| Environmental temperature (°C) | Position-dependent effect | **63419.33** | 63843.76 | 424.43 | 430.43 | 3 | <0.001 | ∗ |
| | Absolute effect | **63419.33** | 63934.36 | 515.03 (90.6) | 523.03 | 4 | <0.001 | ∗ |
| Illumination (ln[Lux/100]) | Position-dependent effect | **63419.33** | 67334.28 | 3914.95 | 3921 | 3 | <0.001 | ∗ |
| | Absolute effect | **63419.33** | 74210.83 | 10791.50 (6876.55) | 10799 | 4 | <0.001 | ∗ |
| Hour of day (after 9am) | Position-dependent effect | **63419.33** | 63460.99 | 41.66 | 53.66 | 6 | <0.001 | ∗ |
| | Absolute effect | **63419.33** | 66802.43 | 3383.10 (3341.44) | 3399.10 | 8 | <0.001 | ∗ |
| Intercept | Position-dependent effect | **63419.33** | 63544.48 | 125.15 | 131.15 | 3 | <0.001 | ∗ |
| | Model intercept | | | | | | | - |

**Supplementary Table S6:** The results of model selection testing weather effects on *Coreopsis verticillata*. Comparisons of standing best models and a simpler version where parameters are removed are given for each effect tested in our model selection process. Bold AIC value indicates best performing model of each comparison. AIC is given for both models (note how standing best model AIC matches one of the previous model AICs). Also given is an assessment of model fit, Δdeviance=Δdev. A verdict on each comparison is given: Asterisks '∗' indicate parameters where AIC comparisons indicate inclusion of effects in best model, based on Richards (2008).

| Tested effect | | Standing best model AIC | Simpler model AIC | ΔAIC | Δdev. | df | *p* | Verdict |
|---|---|---|---|---|---|---|---|---|
| Wind speed (kn) | Position-dependent effect | **100100.0** | 100166.3 | 66.3 | 72.22 | 3 | <0.001 | ∗ |
| | Absolute effect | **100100.0** | 100549.3 | 449.3 (383.0) | 457.28 | 4 | <0.001 | ∗ |
| Pressure (ΔhPa) | Position-dependent effect | 100100.0 | **100094.7** | 5.3 | 0.66 | 3 | 0.883 | |
| | Absolute effect | 100094.7 | **100093.3** | 1.4 | 0.57 | 1 | 0.448 | |
| Relative Humidity (%) | Position-dependent effect | 100093.3 | **100089.2** | 4.1 | 1.88 | 3 | 0.597 | |
| | Absolute effect | **100089.2** | 100428.8 | 339.6 | 341.63 | 1 | <0.001 | ∗ |
| Cloud Cover (oktas) | Position-dependent effect | **100089.2** | 100158.1 | 68.9 | 74.91 | 3 | <0.001 | ∗ |
| | Absolute effect | **100089.2** | 100170.9 | 81.7 (12.8) | 89.75 | 4 | <0.001 | ∗ |
| Environmental temperature (°C) | Position-dependent effect | **100089.2** | 100109.8 | 20.6 | 26.67 | 3 | <0.001 | ∗ |
| | Absolute effect | **100089.2** | 103413.7 | 3324.5 (3303.9) | 3332.5 | 4 | <0.001 | ∗ |
| Illumination (ln[Lux/100]) | Position-dependent effect | **100089.2** | 102386.4 | 2297.2 | 2303.2 | 3 | <0.001 | ∗ |
| | Absolute effect | **100089.2** | 116430.0 | 16340.8 (14043.6) | 16349 | 4 | <0.001 | ∗ |
| Hour of day (after 9am) | Position-dependent effect | **100089.2** | 100128.0 | 38.8 | 50.85 | 6 | <0.001 | ∗ |
| | Absolute effect | **100089.2** | 101872.2 | 1783.0 (1744.2) | 1799.1 | 8 | <0.001 | ∗ |
| Intercept | Position-dependent effect | **100089.2** | 100445.0 | 355.8 | 361.89 | 3 | <0.001 | ∗ |
| | Model intercept | | | | | | | - |

**Supplementary Table S7:** The results of model selection testing weather effects on *Geranium psilostemon*. Comparisons of standing best models and a simpler version where parameters are removed are given for each effect tested in our model selection process. Bold AIC value indicates best performing model of each comparison. AIC is given for both models (note how standing best model AIC matches one of the previous model AICs). Also given is an assessment of model fit, Δdeviance=Δdev. A verdict on each comparison is given: Asterisks '∗' indicate parameters where AIC comparisons indicate inclusion of effects in best model, based on Richards (2008); crosses '†' indicate parameters where inclusion of the parameter in the best model improved fit, but AIC was not improved sufficiently for the parameter to be included.

| Tested effect | | Standing best model AIC | Simpler model AIC | ΔAIC | Δdev. | df | *p* | Verdict |
|---|---|---|---|---|---|---|---|---|
| Wind speed (kn) | Position-dependent effect | **59469.83** | 59488.69 | 18.86 | 24.86 | 3 | <0.001 | ∗ |
| | Absolute effect | **59469.83** | 59604.61 | 134.78 (115.92) | 142.78 | 4 | <0.001 | ∗ |
| Pressure (ΔhPa) | Position-dependent effect | 59469.83 | **59473.26** | 3.43 | 9.43 | 3 | 0.024 | † |
| | Absolute effect | 59473.26 | **59475.44** | 2.18 | 4.18 | 1 | 0.041 | † |
| Relative Humidity (%) | Position-dependent effect | 59475.44 | **59474.25** | 1.19 | 4.81 | 3 | 0.186 | |
| | Absolute effect | **59474.25** | 60149.36 | 675.11 | 677.11 | 1 | <0.001 | ∗ |
| Cloud Cover (oktas) | Position-dependent effect | 59474.25 | **59472.25** | 2.00 | 4.00 | 3 | 0.262 | |
| | Absolute effect | **59472.25** | 59491.50 | 19.25 | 21.25 | 1 | <0.001 | ∗ |
| Environmental temperature (°C) | Position-dependent effect | **59472.25** | 59504.45 | 32.20 | 38.20 | 3 | <0.001 | ∗ |
| | Absolute effect | **59472.25** | 62135.00 | 2662.75 (2630.55) | 2670.7 | 4 | <0.001 | ∗ |
| Illumination (ln[Lux/100]) | Position-dependent effect | **59472.25** | 60151.65 | 679.40 | 685.4 | 3 | <0.001 | ∗ |
| | Absolute effect | **59472.25** | 69988.05 | 10515.80 (9836.40) | 10524 | 4 | <0.001 | ∗ |
| Hour of day (after 9am) | Position-dependent effect | **59472.25** | 59487.89 | 15.64 | 27.64 | 6 | <0.001 | ∗ |
| | Absolute effect | **59472.25** | 60411.80 | 939.55 (923.91) | 955.55 | 8 | <0.001 | ∗ |
| Intercept | Position-dependent effect | **59472.25** | 59728.49 | 256.24 | 262.24 | 3 | <0.001 | ∗ |
| | Model intercept | | | | | | | - |

**Supplementary Table S8:** The results of AIC comparisons testing alternative groupings of flower position effects. For each alternative position model (including the full or 'standing best' model from the previous comparisons) the model's grouping of flower position is given (shared letters between positions indicates that the model treats these positions as equivalent). Also given are the AIC results (df= degree of freedom, delta= difference in AIC from best performing model) for each of the alternative position models. Bold models indicate those selected as best fitting for each species, based on Richards (2008).

| | A) repro. struc. | B) petal base | C) petal Mid. | D) petal tip | *Cistus* 'snow fire' | | | *Cistus* 'snow white' | | | *Coreopsis verticillata* | | | *Geranium psilostomon* | | |
|---|---|---|---|---|---|---|---|---|---|---|---|---|---|---|---|---|
| | | | | | df | AIC | delta | df | AIC | delta | df | AIC | delta | df | AIC | delta |
| Absent | A | A | A | A | 11 | 107215.68 | -21377.17 | 11 | 76762.21 | -13342.88 | 10 | 118251.2 | -18162.00 | 10 | 63355.07 | -3882.82 |
| ap1 | A | B | A | A | 18 | 98533.71 | -12695.20 | 18 | 75030.43 | -11611.10 | 17 | 118237.4 | -18148.23 | 16 | 60528.16 | -1055.91 |
| ap2 | A | A | B | A | 18 | 104735.38 | -18896.88 | 18 | 74824.33 | -11405.01 | 17 | 116379.0 | -16289.83 | 16 | 63286.82 | -3814.57 |
| ap3 | A | A | A | B | 18 | 99292.53 | -13454.02 | 18 | 71807.08 | -8387.75 | 17 | 113335.7 | -13246.50 | 16 | 60976.30 | -1504.05 |
| ap4 | A | B | B | A | 18 | 106216.34 | -20377.83 | 18 | 76738.34 | -13319.01 | 17 | 117071.5 | -16982.38 | 16 | 61653.30 | -2181.05 |
| ap5 | A | B | C | A | 25 | 97843.78 | -12005.28 | 25 | 73918.66 | -10499.33 | 24 | 116232.4 | -16143.27 | 22 | 60218.83 | -746.58 |
| ap6 | A | B | A | B | 18 | 107217.95 | -21379.45 | 18 | 76255.60 | -12836.27 | 17 | 115057.4 | -14968.27 | 16 | 63352.01 | -3879.76 |
| ap7 | A | B | A | C | 25 | 93188.95 | -7350.44 | 25 | 71259.97 | -7840.64 | 24 | 112856.2 | -12767.08 | 22 | 59634.37 | -162.12 |
| ap8 | A | A | B | B | 18 | 88306.41 | -2467.90 | 18 | 64515.01 | -1095.68 | 17 | 107266.1 | -7176.99 | 16 | 60195.06 | -722.81 |
| ap9 | A | A | B | C | 25 | 87321.68 | -1483.17 | 25 | 64131.05 | -711.72 | 24 | 106927.5 | -6838.33 | 22 | 59565.01 | -92.76 |
| ap10 | A | B | B | B | 18 | 105057.73 | -19219.22 | 18 | 71285.75 | -7866.42 | 17 | 103651.8 | -3562.60 | 16 | 63273.73 | -3801.48 |
| ap11 | A | B | C | B | 25 | 103638.48 | -17799.98 | 25 | 70662.86 | -7243.54 | 24 | 103559.3 | -3470.12 | 22 | 63230.41 | -3758.16 |
| ap12 | A | B | B | C | 25 | 98755.40 | -12916.90 | 25 | 68209.12 | -4789.79 | 24 | 101614.3 | -1525.13 | 22 | 60982.21 | -1509.96 |
| ap13 | A | B | C | C | 25 | 86891.51 | -1053.01 | 25 | 63819.45 | -400.12 | 24 | 100529.7 | -440.56 | 22 | 60106.96 | -634.71 |
| **Standing best** | **A** | **B** | **C** | **D** | **32** | **85838.50** | **0.00** | **32** | **63419.33** | **0.00** | **31** | **100089.2** | **0.00** | **28** | **59472.25** | **0.00** |

**Supplementary Table S9:** Summary of the parameter values of the best fitting floral temperature models of each flower species. The values and SEM of each model parameter is given for each species, 'n/a' indicates that the parameter is not included in the best fitting model. To aid parameter identification parameters are named using both the identifiers used in supplementary information 4, column 'parameter' and names given in the generation of these values at the end of the accompanying code for this publication (Harrap et al., 2024), column 'R identifier'. Right aligned R identifiers and parameter names indicates that these pertain to position-dependent effects. R identifiers and parameter names beginning with 'B.' or with subscript 'b' respectively, indicate the position-dependent effect, relative to that of the reproductive structures, of the variable on the petal base. R identifiers and parameter names beginning with 'C.' or with subscript 'c' respectively, indicate the position-dependent effect, relative to that of the reproductive structures, of the variable on the petal middle. R identifiers and parameter names beginning with 'D.' or with subscript 'd' respectively, indicate the position-dependent effect, relative to that of the reproductive structures, of the variable on the petal edge. For weather variables where position-dependent effects are present the base effect of each weather variable described the effects of that variable on the reproductive structures. Where there is no position-dependent effects for a variable the base effect of that variable describes the effects at all positions (see Supplementary Information 4). Additionally, a summary of weather variables experienced across all sampling periods are provided. For variables measured hourly (all variables except those pertaining to illumination, Lux/100), all summary values for these variables are calculated across all hours where sampling took place. In the case of parameters measured at the thermography incident scale (illumination, Lux/100), summary values are calculated across all thermography incidents. Where parameters of the model act on a transformed value of the weather variable, the transformed value parameters act on is given in brackets below the raw values.

| Weather Variable | | | Range of weather variable values | | | | | | *Cistus* 'snow fire' | | *Cistus* 'snow white' | | *Coreopsis verticillata* | | *Geranium psilostemon* | |
|---|---|---|---|---|---|---|---|---|---|---|---|---|---|---|---|---|
| **R identifier** | **parameter** | **units** | **min** | **max** | **median** | **mean** | **1stQ** | **3rdQ** | **Value** | **SEM** | **Value** | **SEM** | **Value** | **SEM** | **Value** | **SEM** |
| Intercept | $i_a$ | | - | - | - | - | - | - | -5.832 | 0.723 | -4.183 | 0.830 | -21.184 | 0.453 | -24.60 | 0.906 |
| B.PetalBase | $i_b$ | | | | | | | | -3.182 | 0.526 | 3.371 | 0.499 | 4.797 | 0.442 | -3.632 | 0.886 |
| C.PetalMid | $i_c$ | | | | | | | | 4.241 | 0.526 | 4.989 | 0.499 | 6.397 | 0.442 | 1.072 | 0.886 |
| D.PetalTip | $i_d$ | | | | | | | | 6.146 | 0.526 | 4.922 | 0.499 | 7.964 | 0.442 | 4.650 | 0.886 |
| HOUR | $h_a$ | h | 9 | 18 | 12 | 12.77 | 11 | 15 | 0.718 | 0.040 | 1.020 | 0.035 | 0.632 | 0.033 | 0.570 | 0.105 |
| B.HOUR | $h_b$ | (h-9) | (0) | (9) | (3) | (3.77) | (2) | (6) | -0.162 | 0.051 | 0.003 | 0.045 | 0.025 | 0.045 | 0.105 | 0.114 |
| C.HOUR | $h_c$ | | | | | | | | 0.230 | 0.051 | 0.120 | 0.045 | -0.099 | 0.045 | 0.068 | 0.114 |
| D.HOUR | $h_d$ | | | | | | | | 0.392 | 0.051 | 0.200 | 0.045 | -0.227 | 0.045 | 0.012 | 0.114 |
| HOUR2 | $j_a$ | h | 9 | 18 | 12 | 12.77 | 11 | 15 | -0.043 | 0.005 | -0.085 | 0.005 | -0.052 | 0.004 | -0.048 | 0.013 |
| B.HOUR2 | $j_b$ | $(h-9)^2$ | (0) | (81) | (9) | (14.21) | (4) | (36) | 0.016 | 0.007 | 0.000 | 0.006 | -0.002 | 0.005 | -0.015 | 0.014 |
| C.HOUR2 | $j_c$ | | | | | | | | -0.019 | 0.007 | -0.009 | 0.006 | 0.014 | 0.005 | -0.005 | 0.014 |
| D.HOUR2 | $j_d$ | | | | | | | | -0.034 | 0.007 | -0.017 | 0.006 | 0.029 | 0.005 | 0.005 | 0.014 |
| LnLux | $l_a$ | lux/100 | 27 | 1401 | 490 | 593.8 | 293 | 897 | 3.555 | 0.037 | 3.365 | 0.032 | 4.493 | 0.037 | 4.789 | 0.124 |
| B.LnLux | $l_b$ | (ln(lux/100)) | (3.30) | (7.24) | (6.19) | (6.39) | (5.68) | (6.80) | 0.744 | 0.042 | -0.457 | 0.038 | -1.246 | 0.049 | 0.701 | 0.134 |
| C.LnLux | $l_c$ | | | | | | | | -1.811 | 0.042 | -1.796 | 0.038 | -1.832 | 0.049 | -0.539 | 0.134 |
| D.LnLux | $l_d$ | | | | | | | | -2.468 | 0.042 | -2.133 | 0.038 | -2.256 | 0.049 | -1.320 | 0.134 |
| Tenv | $t_a$ | °c | 13.9 | 30.4 | 17.7 | 18.26 | 16.5 | 18.93 | 0.251 | 0.024 | 0.263 | 0.028 | 0.798 | 0.018 | 0.762 | 0.026 |
| B.Tenv | $t_b$ | | | | | | | | -0.001 | 0.012 | -0.063 | 0.011 | 0.005 | 0.019 | -0.017 | 0.025 |
| C.Tenv | $t_c$ | | | | | | | | 0.156 | 0.012 | 0.106 | 0.011 | 0.048 | 0.019 | 0.052 | 0.025 |
| D.Tenv | $t_d$ | | | | | | | | 0.179 | 0.012 | 0.142 | 0.011 | 0.084 | 0.019 | 0.063 | 0.025 |
| CloudCov | $o_a$ | oktas | 0 | 8 | 5 | 4.209 | 0 | 7 | 0.042 | 0.007 | -0.063 | 0.007 | -0.084 | 0.009 | -0.039 | 0.008 |
| B.CloudCov | $o_b$ | | | | | | | | n/a | n/a | n/a | n/a | 0.065 | 0.012 | n/a | n/a |
| C.CloudCov | $o_c$ | | | | | | | | n/a | n/a | n/a | n/a | 0.091 | 0.012 | n/a | n/a |
| D.CloudCov | $o_d$ | | | | | | | | n/a | n/a | n/a | n/a | 0.091 | 0.012 | n/a | n/a |
| RH | $r_a$ | % | 47.3 | 100 | 73.9 | 72.49 | 64.7 | 79.45 | 0.028 | 0.005 | 0.024 | 0.005 | 0.044 | 0.002 | 0.064 | 0.002 |
| B.RH | $r_b$ | | | | | | | | 0.001 | 0.004 | -0.000 | 0.004 | n/a | n/a | n/a | n/a |
| C.RH | $r_c$ | | | | | | | | 0.008 | 0.004 | 0.014 | 0.004 | n/a | n/a | n/a | n/a |
| D.RH | $r_d$ | | | | | | | | 0.012 | 0.004 | 0.024 | 0.004 | n/a | n/a | n/a | n/a |
| HLhPA | $p_a$ | hPa | 1005 | 1027 | 1019 | 1018 | 1015 | 1022 | -0.134 | 0.019 | -0.125 | 0.018 | n/a | n/a | n/a | n/a |
| B.HLhPA | $p_b$ | (ΔhPa) | (-8) | (14) | (6) | (5) | (2) | (9) | n/a | n/a | n/a | n/a | n/a | n/a | n/a | n/a |
| C.HLhPA | $p_c$ | | | | | | | | n/a | n/a | n/a | n/a | n/a | n/a | n/a | n/a |
| D.HLhPA | $p_d$ | | | | | | | | n/a | n/a | n/a | n/a | n/a | n/a | n/a | n/a |
| Wind.Sp | $w_a$ | kn | 2 | 24 | 11 | 11.51 | 8 | 15 | -0.098 | 0.007 | -0.145 | 0.007 | -0.096 | 0.005 | -0.101 | 0.018 |
| B.Wind.Sp | $w_b$ | | | | | | | | -0.010 | 0.007 | -0.004 | 0.007 | 0.027 | 0.007 | 0.002 | 0.019 |
| C.Wind.Sp | $w_c$ | | | | | | | | 0.060 | 0.007 | 0.052 | 0.007 | 0.048 | 0.007 | 0.035 | 0.019 |
| D.Wind.Sp | $w_d$ | | | | | | | | 0.087 | 0.007 | 0.068 | 0.007 | 0.051 | 0.007 | 0.058 | 0.019 |
